# Unraveling the impact of host genetics and factors on the urinary microbiome in a young population

**DOI:** 10.1101/2024.05.17.594782

**Authors:** Leying Zou, Zhe Zhang, Junhong Chen, Ruijin Guo, Xin Tong, Yanmei Ju, Haorong Lu, Huanming Yang, Jian Wang, Yang Zong, Xun Xu, Xin Jin, Liang Xiao, Huijue Jia, Tao Zhang, Xiaomin Liu

**Affiliations:** BGI Research, Wuhan 430074, China; BGI Research, Shenzhen 518083, China; College of Life Sciences, University of Chinese Academy of Sciences, Beijing 100049, China; China National Genebank, BGI-Shenzhen, Shenzhen 518120, China; James D. Watson Institute of Genome Sciences, Hangzhou 310058, China; Shenzhen Engineering Laboratory of Detection and Intervention of Human Intestinal Microbiome, BGI-Shenzhen, Shenzhen 518083, China; Institute of Precision Medicine–Greater Bay Area (Guangzhou), Fudan University, Guangzhou, China; Shenzhen Key Laboratory of Human Commensal Microorganisms and Health Research, BGI Research, Shenzhen 518083, China

**Author notes:** These authors contributed equally (Leying Zou, Zhe Zhang, Junhong Chen). To whom correspondence should be addressed: X.L.,; T.Z.,; and.

## Abstract

The significance of the urinary microbiome in maintaining health and contributing to disease development is increasingly recognized. However, a comprehensive understanding of this microbiome and its influencing factors remains elusive. Utilizing whole metagenomic and whole-genome sequencing, along with detailed metadata, we characterized the urinary microbiome and its influencing factors in a cohort of 1579 Chinese individuals. Our findings unveil the distinctiveness of the urinary microbiome from other four body sites, delineating five unique urotypes dominated by *Gardnerella vaginalis, Sphingobium fluviale, Lactobacillus iners, Variovorax sp PDC80* and *Acinetobacter junii*, respectively. We identified 108 host factors significantly influencing the urinary microbiome, collectively explaining 12.92% of the variance in microbial composition. Notably, gender-related factors, including sex hormones, emerged as key determinants in defining urotype groups, microbial composition and pathways, with the urinary microbiome exhibiting strong predictive ability for gender (AUC=0.843). Furthermore, we discovered 43 genome-wide significant associations between host genetic loci and specific urinary bacteria, *Acinetobacter* in particular, linked to 8 host loci (*p* < 5×10^-8^). These associations were also modulated by gender and sex hormone levels. In summary, our study provides novel insights into the impact of host genetics and other factors on the urinary microbiome, shedding light on its implications for host health and disease.

## Introduction

Mounting evidence has implicated the presence of a microbiome community within the urinary tract, commonly referred to as urinary microbiome or urobiome^1,2^. This discovery has been substantiated through high-throughput 16S rRNA/metagenomic sequencing, as well as enhanced urine culture techniques across multiple studies^3–6^. In recent years, the urinary microbiota has garnered considerable attention owning to its potential implications for urinary tract health and diseases. For instance, studies have shown that the urinary microbiome differs significantly in urinary incontinence patients with increased *Gardnerella* and decreased *Lactobacillus* compared to healthy controls^3,7,8^ The urinary microbiome’s composition can also influence susceptibility to urinary tract infections (UTIs) ^9–11^. It has been observed that the postoperative UTIs risk is associated with enrichment of a mixture of uropathogens, as well as depletion of *Lactobacillus* in the preoperative urinary microbiome^12^. Furthermore, the urinary microbiome was shown to exhibit distinct signatures between bladder cancer patients and healthy controls, such as increased abundance of *Fusobacterium* in the patient group^13,14^. Such observations, coupled with the fact that intravesical BCG administration as the gold-standard adjuvant immunotherapy for treating high-risk non-muscle-invasive bladder cancer^15^, suggests a link between urinary microbiome and cancer development and progression potentially through mechanisms involving inflammation and modulation of the immune response.

Thus, better understanding the urinary microbiome and elucidating the factors that influence its composition and diversity are essential for developing effective diagnostic and therapeutic interventions. Previous studies have identified certain host factors, such as sex and age, that may influence composition of the urinary microbiota^16,17^. A recent study in mainly postmenopausal women has suggested a role for host genetics in microbial variations among individuals^18^. However, these studies have predominantly utilized 16S rRNA sequencing that had relatively low resolution and allowed for the identification and quantification of bacterial taxa at the genus level.

Furthermore, existing research has primarily focused on elderly or middle-aged individuals, in which the microbiome may be largely influenced by the environmental factors in the aging process. Thus, there is a gap in knowledge concerning urinary microbiota in young individuals, which our study aims to address. Moreover, there is a necessity to explore additional potential host factors, such as blood metabolites, urine factors, lifestyle, etc., as they may also impact the urinary microbiota. Current research in this area remains limited and calls for more in-depth exploration.

Here in this study, leveraging shotgun metagenomic sequencing and comprehensive metadata from 1579 Chinese individuals from the 4D-SZ cohort^19–24^, which is primarily comprised of young adults around 30 years old, we conducted a thorough investigation to enhance our understanding of the urinary microbiota and its influencing factors. Initially, we characterized the urinary microbiome by analyzing its microbial composition and comparing it to microbial communities in other body sites. We then systematically investigated its influencing factors among 386 traits, encompassing sex, age, BMI, diets, lifestyles, as well as blood and urinary measurements. Moreover, by performing the so far first urinary Metagenome-genome wide association study (M-GWAS) in a subset of 687 individuals with whole genome sequencing data available, we endeavored to unravel the intricate interplay between host genetics and the urinary microbiome.

## Results

### Urinary microbiome: A unique microecology distinct from other body sites

We collected urine samples of 1579 individuals, with 66.12% being females and an average age of 29 years old. Utilizing shotgun metagenomic sequencing, we generated an average of 19.18 ± 7.90 Gb data per sample (**Supplementary Table 1**; **Supplementary Fig. 1**). These samples are from the 4D-SZ cohort^19–24^, with multi-omics data collected including human whole genome, whole metagenomes across multiple body sites, blood metabolites, and detailed questionaries. Since we have metagenomic data from not only the urinary tract but also the other body sites (**Fig. 1a**), including the gut (n=1754), saliva (n=3222), and reproductive tract (n=686), it prompted us to compare the urinary microbiome with the microbial communities inhabiting these distinct body sites. Permutational mutivariate analysis of variance (PERMANOVA) revealed that the urinary microbiome are significantly distinct from the other examined microbiomes. It showed a much lower α-diversity compared to the rich and diverse microbiome harbored in the gut and the oral^25^, with the highest disparity observed in the phylum level with the gut microbiome (R^2^=0.455, a higher R^2^ values corresponding to a larger difference between the compared sites, *p*<0.001; **Supplementary Fig. 2**) and in the genus level with the saliva microbiome (R^2^ = 0.361, *p*<0.001; **Fig. 1b**). In comparison to the vaginal microbiome, which is known to characterize a very low α-diversity^25^, the urinary microbiome exhibited the smallest but significant dissimilarity at both phylum (R^2^=0.214, *p*<0.001) and genus level (R^2^ = 0.111, *p*<0.001) (**Supplementary Fig. 2 and Fig. 1b)**, albeit with a significantly higher α-diversity (**Supplementary Fig. 3)**.

**Figure 1.**
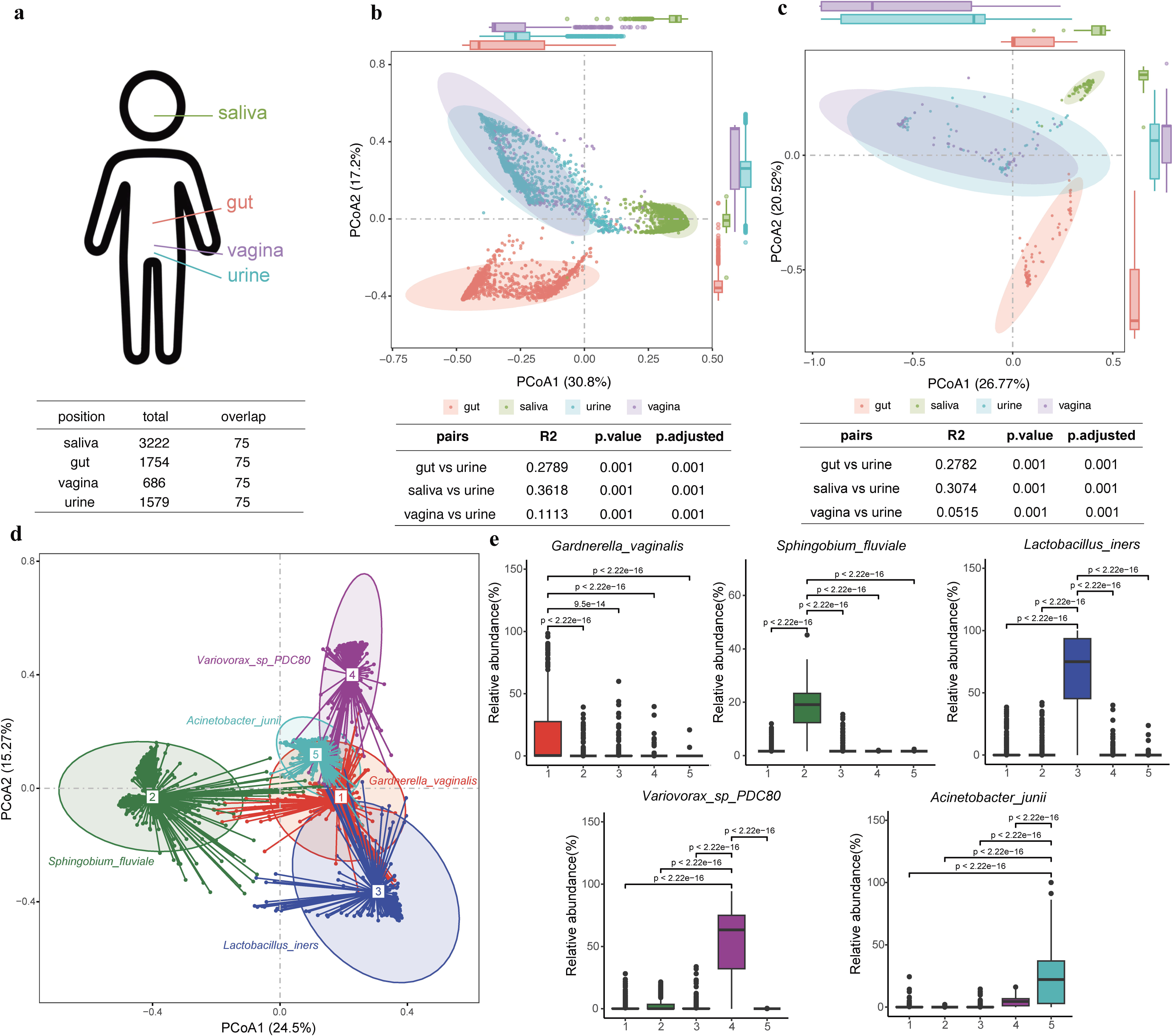
Comparative analysis of microbial communities across human body sites and characterizing the urinary microbiome. **a**) An illustrative diagram presenting the total and shared sample counts of microbial communities from four body sites: saliva, gut, urine, and the reproductive tract. **b**) A scatterplot illustrating the genus-level PCoA distribution of the total microbial samples from saliva, gut, urine, and reproductive tract, along with PERMANOVA pairwise difference values. **c**) A scatterplot showing the genus-level PCoA distribution of 75 shared samples among saliva, gut, urine, and reproductive tract, with PERMANOVA pairwise difference values. **d**) A scatterplot reflecting the species-level PCoA distribution of urinary microbes, the urotypes classification, and the dominant bacteria present in each urotype. **e**) Boxplots displaying the average relative abundance of the five dominant bacteria among individuals with five different urotypes, along with t-test differential analysis.

Further analysis was conducted on metagenomic data from 75 individuals to corroborate the above findings, wherein data from all four body sites were available. Consistent with the broader dataset, the urinary microbiome exhibited the most pronounced differences from the gut microbiome at the phylum level (R^2^ = 0.299; **Supplementary Fig. 4**), and from the salivary microbiome at the genus level (R^2^ = 0.307; **Fig. 1c**), while displaying relatively smaller disparities with the vaginal microbiome (R^2^ = 0.11 at the phylum level and R^2^ = 0.051 at the genus level; **Fig. 1c** and **Supplementary Fig. 4**). Importantly, these variations remained statistically significant (adjusted p < 0.001). Furthermore, we conducted a fast expectation-maximization for microbial source tracking (FEAST^26^) analysis to estimate the potential attributions for the urinary microbiome from the other body sites in these 75 individuals. The results showed the vaginal microbiome was predicted as the main contributor accounting for an average of 60.45% of the urinary microbiome, while saliva (0.48%) and gut (0.27%) microbiome were estimated to have limited contribution in these young adults (**Supplementary Fig. 5**).

It is evident that these distinctions arise from variations in genus composition across different body sites (**Supplementary Table 2**). For example, *Lactobacillus* was the most abundant genus found in both urinary (an average relative abundance of 17.6%) and vaginal samples (58.9%). while its average relative abundance in saliva and gut was less than 0.13%. The second and third most prevalent genera in urine were *Variovora*x and *Acinetobacter*, with mean relative abundances of 10.6% and 9.4%, respectively; however, their average relative abundances in the other examined sites were less than 0.03%. *Prevotella* is found to be enriched in the saliva with an average relative abundance of 29.1%, while it is less abundant in the gut at 13.4%, followed by urine at 6.7% and vagina at 3.6%. The distinctive microbial compositions and species’ diversities across different human body sites resulted in marked dissimilarities in the microbiota across these regions.

In summary, our findings underscore the uniqueness of the urinary microbiota in comparison to other anatomical regions such as the gut, saliva, and vagina. This distinct microbial ecosystem warrants further in-depth investigation.

### Characteristics of the urinary microbiome

Since the urinary microbiome represents a distinctive microecology, we further characterized its community composition in detail. First, we examined the top ten most abundant genera and species in the cohort (**Supplementary Fig. 6**). At the genus level, *Lactobacillus* had the highest relative abundance, followed by *Variovorax, Acinetobacter, Prevotella,* and *Sphingobium*. At the species level, the top five bacteria with the highest relative abundances, in descending order, were *Lactobacillus iners, Variovorax sp. PDC80, Sphingobium fluviale, Lactobacillus crispatus*, and *Gardnerella vaginalis*. Notably, we observed high Pearson correlations between the relative abundances of bacteria commonly found in the vagina, such as *Lactobacillus*, *Gardnerella*, *Prevotella*, and *Ureaplasma*, in the urinary and vaginal microbiomes (Pearson’s r >0.5).

Next, using the partitioning around medoids (PAM) clustering method^27^, the urinary microbial communities were clustered into five distinct clusters (referred to as “urotypes” analogous to the “enterotypes”^28^). By applying Linear Discriminant Analysis (LDA) effect size (LEfSe)^29^ for detecting species discriminating each urotype from the rest and ranking them by LDA score, we determined the dominant species for each urotype. They are *Gardnerella vaginalis* (accounting for 23.10% of the samples)*, Sphingobium fluviale* (34.11%)*, Lactobacillus iners* (15.76%), *Variovorax sp. PDC80* (16.58%), and *Acinetobacter junii* (10.38%), respectively (**Fig. 1d**). Consistent with the clustering results, each urotype showed a significant enrichment of the respective dominant bacteria (**Fig. 1e**).

While the dominant species characterized the highest LDA score for each urotype, there were urotype-characteristic species identified, especially in clusters 1, 2, and 5 **(Supplementary Fig. 7**; **Supplementary Table 3**). For cluster 1 (*Gardnerella vaginalis* urotype), the second-ranked species *Lactobacillus crispatus* followed the dominant bacterium *Gardnerella vaginalis* so close in LDA score, just like their close relative abundances with an average of 3.91% versus 3.92%. The individuals characterizing this urotype also harbored significantly higher levels of species belonging to the genus *Prevotella*, including *Prevotella bivia* (3.67%), *Prevotella timonensis* (3.43%), *Prevotella disiens* (3.08%), *Prevotella colorans* (2.84%), and *Prevotella corporis* (2.70%). For cluster 2 (*Sphingobium fluviale* urotype), species such as *Sphingomonas ursincola* (3.76%), *Novosphingobium sp AAP93* (3.62%), *Erythrobacter donghaensis* (3.59%) and *Acinetobacter johnsonli* (3.56%) were also highly represented. For cluster 3 (*Lactobacillus iners* urotype), *Lactobacillus jensenii* ranked second with an LDA score of 2.7 and belonged to the same genus *Lactobacillus* as the dominant species. In cluster 4 (*Variovorax sp PDC80* urotype), *Bradyrhizobium sp CCH1 B1* also showed a high proportion with an LDA score of 3.76. Finally, in cluster 5 (*Acinetobacter junii* urotype), *Herbaspirillum aquaticum* (3.50%), *Cutibacterium acnes* (3.32%), and *Moraxella osloensis* (3.31%) were also most abundant and significantly enriched compared to the other clusters.

Microbial pathways were also been investigated **(Supplementary Fig. 8; Supplementary Table 4)**. S-adenosyl-L-methionine salvage I (PWY-6151), sucrose biosynthesis II (PWY-7238) and guanosine ribonucleotides de novo biosynthesis (PWY-7221) were enriched in cluster 1 (*Gardnerella vaginalis* as the dominant bacteria). Consistently, *G. vaginalis* contributed to PWY-6151^30^ which has previously been linked to anti-proliferative, pro-apoptotic and anti-metastatic processes in several types of cancer^31^. Similarly, *L. iners* and its contributed pathway pyruvate fermentation to acetate and lactate (PWY-5100, P41-PWY)^30^ were both most enriched in individuals of cluster 3. PWY-5136: Fatty acid beta-oxidation II (plant peroxisome), PWY-7663: gondoate biosynthesis (anaerobic) and PWY-4984: urea cycle were mainly enriched pathways in cluster 1, 4, and 5, respectively. We found bacteria and their contributing pathways were consistently enriched in each urotype cluster.

### Gender-related factors determined urotypes and urinary microbial communities

To understand the factors influencing urotypes, we calculated their correlations with host factors. Notably, gender-related factors emerged as the primary determinants of these urotypes. Menstrual cycle, serum testosterone, serum estradiol, serum creatinine, proportion of urine, menstrual duration, menarche age, hemoglobin, and gender were significantly associated with all five urotypes (**Fig. 2a**, **Supplementary Table 5**). Remarkably, gender distinctly differentiated the urotype groups. Females exhibited higher relative abundances of *Gardnerella vaginalis* (urotype 1) and *Lactobacillus iners* (urotype 3), aligning their urinary microbiome predominantly with these two urotypes. In contrast, males demonstrated higher abundances of *Sphingobium fluviale* (urotype 2)*, Variovorax sp. PDC80* (urotype 4), and *Acinetobacter junii* (urotype 5), characterizing their urinary microbial communities with these three respective urotypes. Additionally, the gender-biased urotypes displayed significant correlations with gender-dependent metadata. For instance, urotypes 4 and 5, predominantly found in males, correlate with higher levels of serum testosterone, urine specific gravity, muscle mass, basal metabolic rate, serum creatinine, and serum uric acid — clinical and biochemical parameters which are typically elevated in males compared to females. Conversely, urotypes 1 and 3, more common in females, show a significant correlation with increased serum estradiol levels. Intriguingly, the gut microbiota also display sex differences, correlating significantly with gonadal steroids such as serum testosterone^32^. Additionally, it is well-established that estrogen can promote vaginal *Lactobacillus* level by accumulated glycogen production in the vaginal squamous epithelial cells^33^. It might be reasonable to speculate that, the relatively higher levels of estrogen in women, which are partly excreted through urine, may similarly also be in relation with enhanced growth of *Lactobacillus* within the urinary tract.

**Figure 2.**
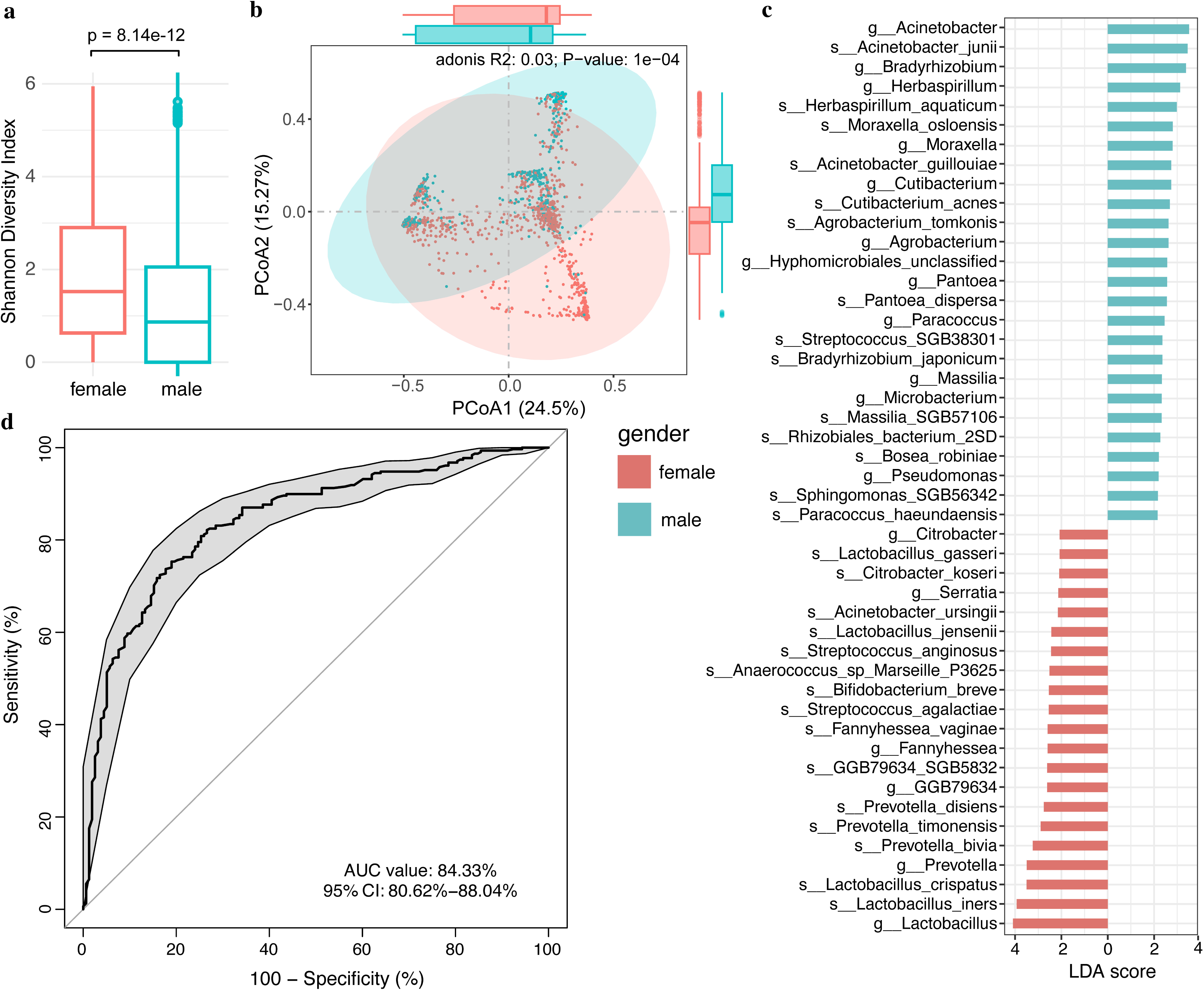
Impact of host phenotypes on urinary microbiota. This scatterplot presents the distribution of Pearson correlation estimate values between host phenotypes and five urotypes, highlighting the top 20 significant associations for each urotype, with colors corresponding to the urine classification results in Figure 1d. Each phenotype is ranked from high to low according to the average of Pearson correlation estimate value for each urotype. **b**) A bar chart showing the top 40 significant host phenotypes sorted by their explained variance (R^2^ values) in the β-diversity of urinary microbiota. **c**) A categorized bar chart displaying the explained variance of significant host phenotypic factors for the urinary microbiota β-diversity, including: 108 total independent host factors, 80 blood factors, 3 urinary factors, gender, 6 anthropometric factors, 6 health factors, 7 lifestyle factors, and 5 diet-related factors.

We further delved into host factors that contributed to the overall composition of the urinary microbiome and identified a total of 137 significant influencing factors (p.adjust < 0.05; **Supplementary Table 6**). Serum testosterone, gender, urine specific gravity, muscle mass, basal metabolic rate, serum creatine and estradiol were among the top influencing factors associated with β-diversity of the urinary microbiome (**Fig. 2b**). Due to some factors were highly correlated, we calculated the Spearman correlations among the 137 host factors and removed 29 collineated factors (Spearman’s r>0.6 || r<-0.6). The remaining 108 independent and significant covariates collectively explained 12.92% of the variance of microbial composition (**Fig. 2c**), which encompassed 80 blood factors (explained R^2^ = 10.57%), three urinary factors (3.80%), gender (3.45%), six anthropometric measurements (3.16%), six health-related factors (1.84%), seven lifestyles (1.19%) and five dietary factors (0.85%). Among the 80 blood factors, hematocrit (2.3%), serum uric acid (2.0%), 3-Methyl-histidine (1.8%) and isoleucine (1.8%) exhibited substantial impacts on the urinary microbiome. The collective urinary factors ranked as the second most significant influence among the seven categories, with urine specific gravity demonstrating the highest explanatory power (3.28%). Among the six anthropometric factors, height (1.83%) and systolic pressure (1.03%) had explanatory powers exceeded BMI (0.76%). Within the health-related factors, the test result of the free total prostate specific antigen had the most significant impact on the urinary microbiome (1.14%). In contrast, dietary factors had a comparatively lower effect, with the highest influencing factor being fat meat favorites, which had an explanatory power of only 0.25% (**Supplementary Table 6).**

In addition to individual urotypes and microbial composition, we also identified 69 host factors significantly associated with 115 microbial pathways (p.adjust < 0.05; **Supplementary Table 7**). Consistent with the findings for microbial composition, urine specific gravity, serum testosterone and gender were the top three strongest host factors correlated with various of microbial pathways. They were all linked to pathways such as HISTSYN-PWY: L-histidine biosynthesis, HEME-BIOSYNTHESIS-II: heme b biosynthesis I (aerobic) HEME-BIOSYNTHESIS-II-1: heme b biosynthesis V (aerobic), PWY-5136: fatty acid beta-oxidation II (plant peroxisome) and TRPSYN-PWY: L-tryptophan biosynthesis. Testosterone administration has been reported to reduce the urinary excretion of histamine in female mice^34^ and increased plasma tryptophan levels in women^35^. Further research is warranted to elucidate the precise mechanisms through which these factors interact with the microbial pathways.

Motivated by the insights that gender and gender-related factors are primary influencers of the urinary microbiome, we proceeded to investigate differences in the microbial community between genders in detail. We first observed significantly higher α-diversity, as measured by the Shannon index, in the urinary microbiota of females than in males (*p* < 8.14×10^-12^, **Fig. 3a**). The microbiota composition also significantly differed between males and females (R^2^ = 0.03, *p* < 1×10^-4^, **Fig. 3b**). Furthermore, we identified 47 sex-differentiated microbial features including 18 genus and 29 species, by using LEfSe analysis (**Fig. 3c**). Females were characterized by higher abundances of genera *Lactobacillus*, *Prevotella, Fannyhessea, Serratia*, and *Citrobacter*. However, males exhibited higher abundances in species belonging to *Acinetobacter, Bradyrhizobium, Herbaspirillum, Moraxella*, *Cutibacterium*, and other genera. These results were in line with the Spearman correlations between bacterial profiles and serum testosterone (**Supplementary Fig. 9**), suggesting the important effects of sex and sex hormones on shaping the urinary microbiota.

**Figure 3.**
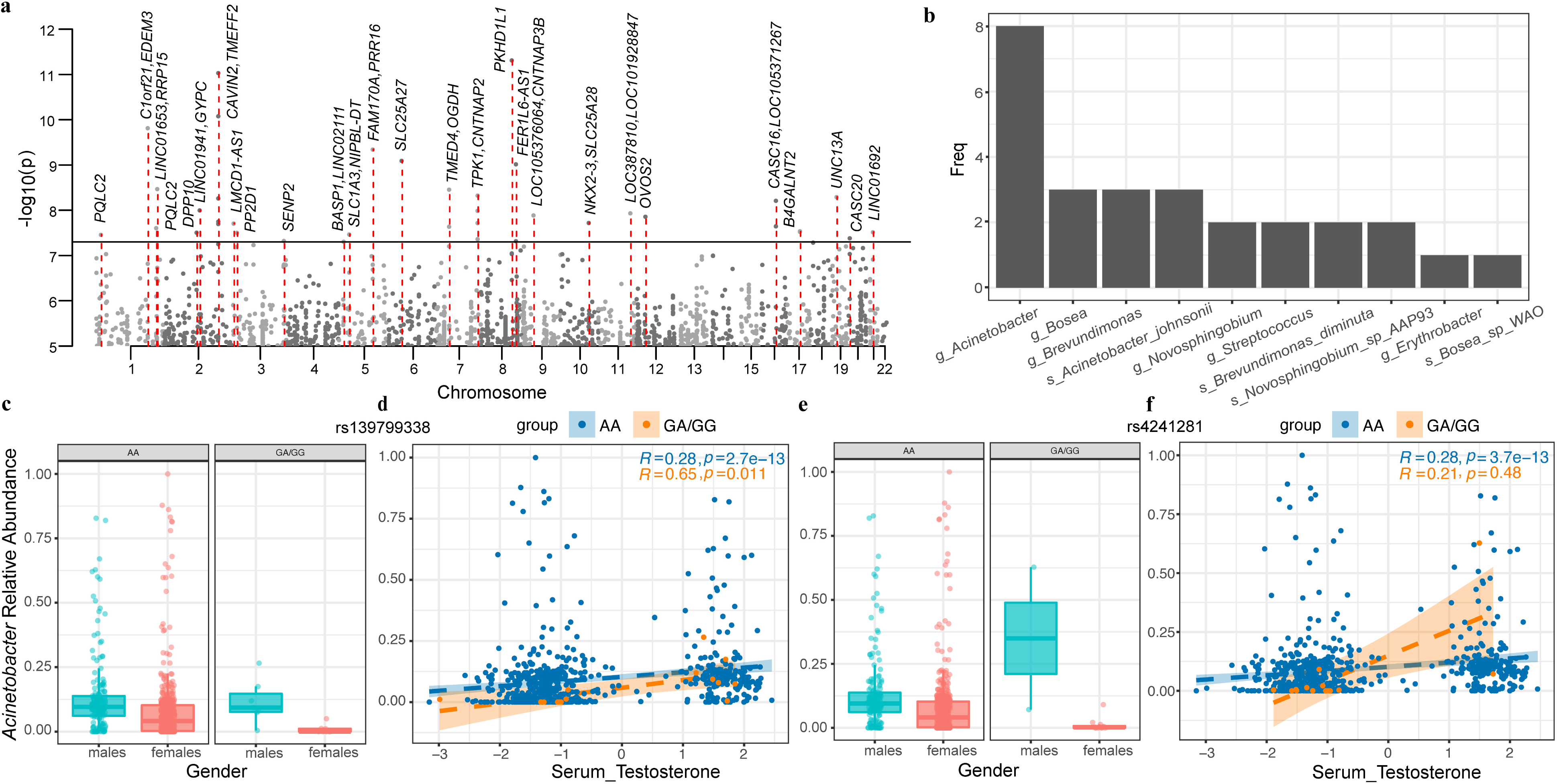
The urinary microbiome differed between genders and effectively distinguished gender. **a)** Box plot of α-diversity (Shannon index) of the urinary microbiome between males and females. **b)** β-diversity of the urinary microbiome using principal coordinate analysis, as colored by males (cyan) and females (red). The first and second axes, PCoA1 and PCoA2, explaining the highest variance, are shown. Dashed ellipses represent the 95% confidence level of permutational multivariate analysis of variance (PERMANOVA) test. Explained variance (R^2^) and p-value were also been showed. **c**) The sex differential urinary microbiota, as sorted by LDA scores in the LEfSe analysis**. d**) A ROC curve demonstrating the predictive power of the urinary microbiota for gender using a random forest model. The analysis comprises 520 male samples and 1044 female samples.

Given the striking gender-related difference in the microbial community, we employed a random-forest-based machine learning approach to assess the predictive capability of urinary microbiota for gender classification. We used the 47 sex-differentiated microbial features for random forest modeling, with 70% of the samples as training sets and 30% as test sets. The predictive accuracy demonstrated by the urinary microbiota in determining gender was remarkably high, with an area under the curve (AUC) value of approximately 0.843 (**Fig. 3d**). Notably, the representative species of urotypes including *Lactobacillus iners* and *Acinetobacter junii* were among the top 10 species contributed most significantly to gender differentiation according to Gini importance (**Supplementary Table 8**). These findings highlight the intricate relationship between microbial composition and host biological characteristics, suggesting that the urinary microbiome’s distinct makeup is reflective of and influenced by the host’s biological sex.

### Host genetics’ influences on the urinary microbiota

Out of a total of 1579 individuals, 687 had host whole genome sequencing data available, which enabled us to investigate the influence of the host genome on the urinary microbiome. To achieve this, we conducted a metagenome-genome-wide association study (M-GWAS) on 106 urinary taxa present in over 10% individuals (**Supplementary Table 9**). After adjusting for covariates such as gender, age, BMI, sequencing read counts, and the top ten host Principal Components (PCs), we identified 43 genome-wide significant associations involving 27 host genetic loci and 10 urinary taxa (*p* < 5×10^-8^, **Fig. 4a**, **Supplementary Table 10**).

**Figure 4.**
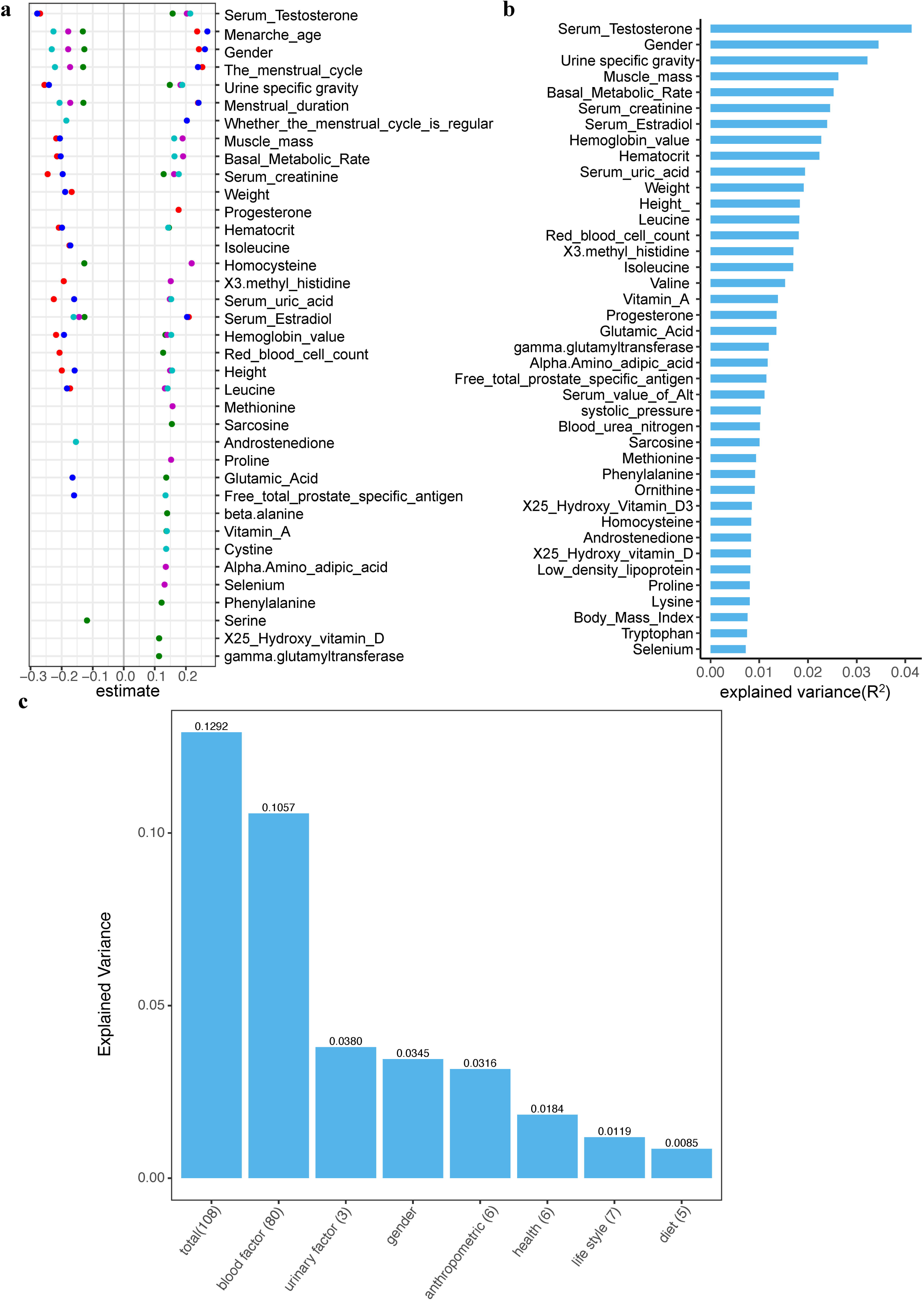
Influences of host genetics and their interactions with gender/sex hormones on urinary microbiota. **a)** A Manhattan plot showing the genetic variants associated with urine microbial taxa (n=106), with genome-wide significance denoted by horizontal black lines (p[=[5[×[10^−8^) and gene names annotated for 27 significant loci. **b)** A bar chart displaying the urinary microbes associated with the significant 27 loci and the number of associations. **c**) Boxplots illustrating the average relative abundance of genus *Acinetobacter* among individuals with different genotypes (AA vs GA/GG) at the top locus rs139799338, as well as stratified by male and female samples. **d**) A scatterplot depicting the association between serum testosterone and the genus *Acinetobacter* relative abundance among individuals with different genotypes (AA vs GA/GG) at the top locus rs139799338. Spearman correlation’s R and p values were also listed. **e**) Boxplots showing the average relative abundance of genus *Acinetobacter* among individuals with different genotypes (AA vs GA/GG) at the top locus rs4241281, as well as stratified by male and female samples. **f**) A scatterplot illustrating the association between serum testosterone and the genus *Acinetobacter* relative abundance among individuals with different genotypes (AA vs GA/GG) at the top locus rs4241281. Spearman correlation’s R and p values were also listed.

Of all the microbial taxa examined at the genus and species level, genus *Acinetobacter* was associated with most host genetic loci (n =8; **Fig. 4b**). The strongest association was observed for SNP rs139799338, located in the intronic region of the *PKHD1L1* gene (*p*=4.82×10^-12^), negatively associated with the relative abundance of the *Acinetobacter* genus (**Fig. 4a**). *PKHD1L1* has been found to be significantly elevated in the urine of clear cell renal cell carcinoma patients^36^. It was also reported to primarily express in both infant and adult human kidneys and is associated with autosomal recessive polycystic kidney disease^37^. The SNP rs139799338 demonstrated associations with various metabolic traits, including the tumor marker carbohydrate antigen 199 (*p*=0.006), glycine (*p*=0.01), vitamin A (*p*=0.01), beta-alanine (*p*=0.03), and cobalt (*p*=0.047), as revealed by the genome-wide association study (GWAS) analysis of host genetic variants and metabolic traits in this cohort (**Supplementary Fig. 10, Supplementary Table 11**).

The second most significant correlation identified was with rs4241281, located in the intergenic region of genes *CAVIN2* and *TMEFF2*, which was found to have a negative association with the genus *Acinetobacter* (*p*=9.13×10^-12^, **Fig. 4a**, **Supplementary Table 10**). This genetic locus was also correlated with the relative abundance of genus *Bosea*, ranking as the third strongest association. Additionally, it was also linked to the abundance of species *Acinetobacter johnsonii* belonging to the genus *Acinetobacter* (*p*=1.75×10^-8^). *CAVIN2* encodes Caveolae Associated Protein 2 in the kidney, and its removal leads to caveolae loss, while overexpression results in caveolae deformation and membrane tubulation^38^. *TMEFF2* has been identified as an epigenetic biomarker for bladder cancer^39^, showing significant associations with increasing tumor grade and stage. Notably, SNP rs4241281 in the *CAVIN2* - *TMEFF2* was also correlated with several blood and urinary factors, including concentrations of mercury (*p*=0.007), alpha amino adipic acid (*p*=0.015), albumin (*p*=0.035), body fat mass and rate(*p*=0.034∼0.017), and urine PH (*p*=0.047**, Supplementary Table 11**). Individuals with the rs185971980 SNP located in the intergenic region of *FAM170A* and *PRR16*, exhibited lower abundance of *Acinetobacter johnsonii* in the urine (*p*=4.46×10^-10^). Additionally, the rs185971980 was correlated with levels of mean hemoglobin content, mean erythrocyte hemoglobin concentration, average RBC volume and blood iron. Similarly, individuals with the rs117738684 SNP had lower abundance of *Acinetobacter johnsonii* (*p*=3.44×10^-9^). The rs117738684 was also correlated with levels of blood testosterone, fasting sugar, methionine, and albumin. Moreover, the SNP rs9975224 within *LINC01692* was associated with not only the abundance of species *Novosphingobium sp. AAP93* but also the urine pH (β=0.087, *p*=9.3×10^-4^). SNPs within *SENP2* was associated with not only the abundance of genus *Acinetobacter* but also the urine specific gravity (β=-0.087, *p*=1.8×10^-4^, **Supplementary Table 11**). These results suggested host genes may influence the urinary microbiota by regulating the blood metabolites and urinary environment.

Gender and sex hormones have demonstrated significant impacts on the urinary microbiome, prompting an exploration of gender-gene interactions using the top two loci associated with specific taxa (**Fig. 4c-f**). An intriguing observation surfaced in our study, revealing that males with heightened testosterone levels displayed an increased abundance of *Acinetobacter* compared to females, regardless of their genetic makeup (genotypes AA or GA/GG at SNP rs139799338). Additionally, carriers of genotype AA exhibited a higher abundance of *Acinetobacter* compared to those carriers of genotype GA/GG in females, while no apparent distinctions were noted among males with varying genotypes (**Fig. 4c**). Similarly, an interactive effect between rs139799338 genotypes and serum testosterone levels on *Acinetobacter* was observed. Specifically, a stronger correlation of serum testosterone level and *Acinetobacter* abundance was noted in individuals with genotype GA/GG (R=0.65, *p*=0.011) compared to individuals with genotype AA (R=0.28, *p*=2.7×10^-13^, **Fig. 4d**). The gender-gene interaction was also evident for another SNP, rs4241281, linked to *Acinetobacter*, with males exhibiting a higher average relative abundance than females (**Fig. 4e**). In the individuals with the rs4241281 AA allele, serum testosterone demonstrated a strong correlation with *Acinetobacter* (R=0.28, *p*=3.7×10^-13^, **Fig. 4e-f**).

### Potential causal relationships between urinary microbiome and diseases

Finally, to explore the potential causal relationships between urinary microbes and diseases by leveraging microbiome-associated host genetic variations, we performed a two-sample Mendelian randomization (MR) analysis. After adjusting for 42 diseases from the Biobank Japan (BBJ) cohort^40^, we identified seven suggestive causal relationships (p<=1.19×10^-3^=0.05/42, **Supplementary Table 12, Supplementary Fig. 11**). The strongest MR evidence suggested a link between type 2 diabetes (T2D) and a decreased abundance of *Streptococcus* SGB38301 (β=-0.076, *p*=2.83×10^-4^). *Paracoccus haematequi* was causally correlated with a decreased risk of T2D (β=-0.013, *p* = 5.12×10^-4^), while the males-enriched *Variovorax sp. PDC80* linked to an increased T2D risk (β=0.017, *p* = 8.89×10^-4^). Genus *Herbaspirillum* potentially increased the risk of atopic dermatitis (β=0.064, *p* = 4.41×10^-4^). *Bosea robiniae* potentially increased the risk of prostate cancer (β=0.058, *p* = 4.65×10^-4^). Keloid was associated with a lower *Bosea* (β = -0.061, *p* = 7.64×10^-4^). These inferred relationships underscore a potential connection between urinary microbes and autoimmune diseases, warranting validation in future studies.

## Discussion

This study represents the initial profiling of the urinary microbiome in young adults in comparison to well-studied human microbiomes from diverse body sites such as the gut, oral cavity, and reproductive tract. While previous studies have also compared the microbial composition across various body sites^41^, they relied on databases from different populations and generally involved smaller sample sizes. In contrast, our research provides a more cohesive and comprehensive analysis by utilizing a uniform cohort.

The urinary microbiome was found to possess unique characteristics compared to other sites, evident in the composition of microbial species, as well as α and β diversity differences. The urinary microbiome exhibited significant dissimilarity from other sites, as demonstrated by PCoA analysis in both the entire sample set and the subset of 75 individuals with data from different sites. The difference in microbial communities across body sites was primarily driven by specific taxa. For instance, the gut microbiome was predominantly composed of the genera *Bacteroides* and *Prevotella*, whereas the saliva microbiome showed a gradient of abundance among *Prevotella, Neisseria*, and *Haemophilus*. The urinary microbiome was predominantly composed of genera such as *Lactobacillus*, *Variovorax*, and *Acinetobacter*, while the vaginal microbiome exhibited more abundant *Lactobacillus* and *Gardnerella*. This highlights the significant role that specific microbial taxa play in characterizing the microbial landscape of different body sites, indicating distinct ecological niches and potentially different functional roles within the human body. Despite the differences, urine and vagina harbored several predominant bacteria in common, supporting the relatively closer relatedness between them^42^ compared to other physically distant sites.

We identified five urotypes and their respective dominant bacteria. A previous study extracted microbiota from encrustations on ureteral stents in patients without urinary tract infections and conducted urotype analysis^43^. Their findings, similar to ours, identified types dominated by microbes such as *Lactobacillus*, *Gardnerella vaginalis*, *Achromobacter*, *Corynebacterium*, and *Streptococcus* (**Supplymentary Table 3**). However, they identified extra bacteria like Actinomycetales, Enterobacterales, and *Staphylococcus* which were less abundant in this cohort. These differences probably have stemmed from the distinct sources of the analyzed microbiome. Their study focused on bacteria from ureteral stents, on which nosocomial bacteria may form biofilm. They employed the 16S rRNA sequencing method, which differs from our metagenomic sequencing approach.

Although some studies indicated the urinary microbiome may be impacted by sex^44–47^, they were performed in small samples and mainly based on the 16S rRNA data, resulting in limited sex-specific bacteria identification. In this study, we discovered extensive sex differences in the urinary microbiota which could well predict sex (AUC=0.843). We also observed a strong link between the urinary microbiome and sex hormone levels as well as some other sex-specific factors. Urinary tract infections (UTIs) are among the most common bacterial infections affecting our urinary system^48^, with uropathogenic *Escherichia coli* being the primary causative agent^11^. Among healthy women aged 18–39 years, *E. coli* was responsible for 80% of UTI cases^49^. In this study, we found that *E. coli* exhibited a higher abundance in women (2.96%) than in men (0.86%). *E. coli* was also enriched in urotype 1, a female-dominated urotype characterized by *Gardnerella vaginalis*. This may help to explain why the UTI incidence was higher in females than in males^48,50^.These findings not only enrich our understanding of the gender-specific susceptibilities to certain urinary tract conditions or diseases but also open avenues for utilizing microbial profiles in gender-specific medical research and potential diagnostic applications.

Our study also marks the first detailed exploration of the impact of blood metabolites on the urinary microbiome. Serum hormones, particularly serum testosterone and serum estradiol, emerged as significant influencers of the urinary microbiome. Other metabolites, such as serum creatinine and serum uric acid, along with blood amino acids like leucine, isoleucine, valine, glutamic acid, and vitamin A, were also found to play crucial roles, highlighting the intricate relationship between various serum components and the composition of the urinary microbiome. Serum creatinine has been regarded as a biomarker of renal function for almost a century^51^ and is widely utilized in clinical diagnostics. In our study, serum creatinine showed significant explanatory power for the β-diversity of urinary microbiota and different urotypes, suggesting that serum creatinine levels might influence kidney function through its effects on the composition of urinary microbiota. The potential causal relationship warrants further investigation. This link emphasizes the intricate connections between metabolic biomarkers and microbiome composition, highlighting the importance of a holistic approach to understanding renal health and disease.

Limited studies investigated the effect of host genome on the urinary microbiome, except a previous study that reported considerable genetic contribution and identified heritable *Escherichia* (A = 0.165) and *Lactobacillus iners* (A = 0.177) by analyzing midstream urine samples from 1,600 older females in the TwinsUK cohort^18^. This study represents a pioneering effort in conducting a genome-wide association analysis of the urinary microbiome along with the host genome. We have successfully identified 27 significant genomic loci associated with 10 urinary microbes, demonstrating the influence of genetic factors on the composition of the urinary microbiome. Genus *Acinetobacter* and its species *Acinetobacter johnsonii* showed the most link to host genetic loci. In addition to the urinary microbiome, *Acinetobacter johnsonii* exhibited a genetic attribute to the skin microbiome^52^. In this GWAS, we also observed an association between genus Acinetobacter and the *MARCO* gene (β=-0.183, *p*=3.67×10^-6^), which encoded macrophage receptor with collagenous structure and correlated with stress urinary incontinence (replication *p*= 0.003) in a GWAS of European women^53^. Moreover, we delved deeper into exploring the impact of interactions between significant loci genotypes, gender, and serum testosterone levels on specific urinary microbes. The insights gained from these analyses offer valuable information on the complex interplay between host genetics, hormonal levels, and microbial abundance.

## Methods

### Study subjects

All the participants involved in this research belonged to the ’4D-SZ’ cohort, which encompassed the collection of blood, gut, saliva, vaginal, and urine samples, along with comprehensive metadata, for a multi-omics investigation as previously reported^19–24^. In this study, 1579 urine samples from the cohort were collected for whole metagenomic sequencing (**Supplementary Table 1**). The detailed urine sampling, sequencing, and quality control design were described below. For comparison, we also included metagenomic samples from the gut (n=1754), saliva (n=3222), and reproductive tract (n=686), all from the 4D-SZ cohort. Feces were collected with an MGIEasy kit, and stool DNA was extracted in accordance with the MetaHIT protocol as described previously^54^. For the salivary sample, we employed a 2× concentration of a stabilizing reagent kit and collected 2 mL of saliva. The DNA extraction from saliva samples was conducted using the MagPure Stool DNA KF Kit B (no. MD5115-02B). For reproductive tract samples, the extraction of genomic DNA from was performed in accordance with the established protocol^55^(Hao L,2020). The DNA concentrations from blood, stool, oral, and reproductive tract samples were estimated by Qubit (Invitrogen), and the library preparation and sequencing were conducted in accordance with the procedures described in previous studies^19–24^.

All recruitment and study procedures involving human subjects were approved by the Institutional Review Boards (IRB) at BGI-Shenzhen (IRB number: 19121). All participants provided written informed consent at enrollment.

### Urine metagenomic sampling, sequencing, and quality control

Midstream voided urine was self-collected using a 40-ml urine cup by 1579 individuals from the 4D-SZ cohort. From the urine cup, 5ml urine was flowed into a sterile tube containing 5ml BGI stabilizing reagent^56^ for the preservation of metagenome at room temperature and then stored at -80°C within the day before subsequent metagenomic analysis.

The samples were thawed and transferred into a 15mL centrifuge tube. Then 1mL TE buffer (1M Tris-HCL & 0.1M EDTA) for crystal dissolution^57^ and 200uL 50mg/mL lysozyme were added before centrifuge at 8000rpm for 10min. Then DNA extraction was proceeded with MagPure Stool DNA KF Kit B (MD5115, Magen)^58^.

DNA extraction and library preparation were performed along with blank negative controls and mock community controls. Specifically, ZymoBIOMICS® Microbial Community Standard (D6300) and ZymoBIOMICS® Microbial Community DNA Standard (D6305) were 1×10^l4 times diluted using saline to simulate the low microbial load in urine samples and divided into aliquots (1ml per tube) for storage at -80°C before using as extraction and library preparation controls respectively. During extraction, blank negative control and extraction control (D6300 diluted) were added for each 96-well plate and processed with the same procedure as for the urine samples. Similarly, separate blank negative control and library preparation control (D6305 diluted) were also added for each 96-well plate during library preparation and processed with the same procedure as for the urine samples. Notably, none of the negative controls for both extraction and library preparation passed the library preparation stage for PCR concentration lower than the minimum criteria of 1ng/uL. In total, 33 extraction mock controls and 43 library mock controls generated metagenomic sequencing data.

### Filtering and Removal of Possible Contaminants

As low biomass microbiome tends to be more easily influenced by contaminants and cross-talk, multiple contamination control steps were further applied to the sequencing data.

Step 1. For all the samples in the entire cohort, species were considered as possible contaminants and collectively removed if a species meets one of the following criteria:

A. Relative abundance significantly correlated with PCR concentration (Spearman test, p-value <0.05)
B. Decontam score <0.3 calculated by ‘decontam’ package^59^ based on relative abundance and PCR concentration or occurrence in <4 samples
C. Detected in the 76 mock controls but other than the 10 expected species comprising the mock community with > 1×10^-4^ mean relative abundance
D. Reads count less than 1000

Step 2. For all the samples in each specific 96-well plate, remaining species other than the 10 expected species detected in the respective mock control with >1×10^-4^ relative abundance were considered as contaminants specific to that 96-well plate and were removed.

Step 3. Filtering out non-bacterial microbes and retaining only bacteria taxa and renormalization

During this process, we found the average fraction of contaminated reads across samples is 4.34% with a median value of 0.45% (**Supplementary Table 1**). Specifically, 68% and 95% of individuals exhibited contaminated fractions below 5% and 15%, respectively (**Supplementary Fig. 12**).

### Constructing the urinary metagenomic taxonomic and functional profile

Taxonomy assignment was performed using MetaPhlAn4^60^ version 4.0.6 with default settings. Compared to its predecessor, MetaPhlAn 3, MetaPhlAn 4 identifies a broader and more detailed range of urinary microbiota. Utilizing this tool, we obtained a raw microbial taxonomic dataset composed of a total of 1,063 taxa (11 phyla, 32 classes, 60 orders, 115 families, 248 genera, and 597 species). The gene functional profiling was performed using HUMAnN v3.8^61^ with databases downloaded from https://huttenhower.sph.harvard.edu/humann_data/. Then, we identified a raw microbial pathway dataset composed of 616 metabolic pathways. At last, we mainly used 208 taxa and 158 pathways with a relative abundance higher than 0.0001 and present in more than 10% of individuals for subsequent analysis.

### Source tracking analysis of urinary microbiome

We used R package ‘FEAST’ to conduct source tracking analysis on 75 samples with microbiome sequencing data of all four body sites, by utilizing gut, saliva, and vaginal microbiota as sources, with urinary microbiota serving as the sink (both in species level). Results were obtained for the percentage contribution of each body site.

### Calculating **α**-, **β**-diversity, PCOA and urotypes

The microbial diversity metrics, including α-diversity (Shannon and Simpson indices) and β-diversity (Bray–Curtis dissimilarities), were calculated based on species-level abundance data using the ’diversity’ and ’vegdist’ functions within the R package ’vegan’, respectively. Subsequently, principal coordinates analysis (PCoA) was conducted on the computed beta-diversity dissimilarities utilizing the ’capscale’ function from the ’vegan’ package.

For sample clustering based on relative species abundances, the JSD distance metric and the Partitioning Around Medoids (PAM) clustering algorithm^27^ were employed. The optimal number of clusters was determined using the Calinski-Harabasz (CH) Index^62^ (**Supplementary Fig. 13**). Following this, the Linear Discriminant Analysis Effect Size (LEfSe)^29^, a nonparametric cross-sample test utilizing Linear Discriminant Analysis (LDA) for test statistic computation, supported by traditional univariate tests for feature selection, was utilized. The microbial taxa and pathways were ranked based on their LDA values in descending order, and the predominant bacteria within each urotype were identified as the species with the highest LDA value.

### Host factors explained the variance of the urinary microbiome

We investigated the correlations between urinary microbiome compositions and 420 host variables encompassing anthropometric measurements, blood and urine parameters, among others. To identify variables significantly linked to β-diversity, we employed Bray-Curtis distance-based redundancy analysis (dbRDA) and determined the variance explained by these factors using the ’capscale’ function in the vegan package. Significance of each response variable was verified through analysis of variance (ANOVA) for the dbRDA using the ’anova.cca()’ function in the vegan package. Only variables showing significant associations (Benjamini-Hochberg FDR < 0.05) with β-diversity in univariable models were included in the multivariable analysis. Due to some factors were highly correlated, we calculated the Spearman correlations among host factors and removed those collineated factors (Spearman’s r>0.6 || r<-0.6). The remaining independent and significant factors were categorized into distinct groups including urinary parameters, anthropometric measures, health-related indicators, lifestyles, and dietary elements. To assess the combined explanatory power of these variable groups on urinary microbiome compositions, we performed a variation partitioning analysis using the vegan package. The ’adj.r.squared’ value was calculated using the RsquareAdj option to quantify the proportion of variance explained by the different categories.

We conducted a Pearson correlation analysis on the 158 metabolic pathways and 420 host factors using the cor.test() function in R (**Supplementary Table 7**).

### High-depth whole-genome sequencing for the blood samples

Among the 1,579 individuals, 687 with blood samples were sequenced to a mean of 30× for the whole genome. For blood samples, buffy coat was isolated and DNA was extracted using HiPure Blood DNA Mini Kit (Magen, catalog no. D3111) according to the manufacturer’s protocol. The library construction and sequencing methods are consistent with those described in previous articles^19^.

For genomic analysis, whole-genome reads were aligned to the GRCh38/hg38 human genome using BWA^63^ (v0.7.15), with low-quality reads filtered out. Alignments were processed into BAM files using Samtools^64^ (v0.1.18) and PCR duplicates were marked with Picardtools (v1.62). The Genome Analysis Toolkit (GATK^65^, v3.8) facilitated SNP and INDEL identification and recalibration, referencing dbSNP (v150). Variant discovery and processing were performed with GATK’s HaplotypeCaller and subsequent tools. The final dataset, comprising 7.2 million SNPs/INDELs(MAF ≥ 1%), were remained for M-GWAS analyses.

For variant inclusion, we set stringent thresholds: (i) a minimum average depth of over 8x; (ii) Hardy-Weinberg equilibrium (HWE) P-value >10−5; and (iii) a genotype calling rate >98%. Regarding sample criteria, we required: (i) mean sequencing depth >20[×; (ii) a variant call rate >98%; (iii) absence of population stratification, confirmed through principal components analysis (PCA) conducted using PLINK^66^(version 1.07); and (iv) the exclusion of related individuals, determined by calculating pairwise identity by descent (IBD, with a Pi-hat threshold of 0.1875) in PLINK. Following these rigorous quality control measures, 687 individuals were left for subsequent analysis.

### Sex-stratified analysis and predictive analysis using urinary microbiota

We used LEfSe analysis to discover sex-differentiated microbiota by applying an LDA threshold of 2.0 and a significance cut-off less than 0.05. Further, we employed the Random Forest algorithm to explore the potential of 47 sex-differentiated microbiota (18 genus and 29 species) for gender prediction. This approach was chosen due to its efficacy in handling complex biological data and its robustness in feature selection, making it particularly suitable for high-dimensional datasets like microbiome compositions. We first divided our dataset into 70% training and 30% testing sets, ensuring a balanced representation of male and female samples. The Random Forest model was then trained on the training set with gender as the response variable and the average relative abundance of different urinary microbes as predictor variables. We used 500 trees in our model to ensure adequate learning and generalization capability. The model’s performance was evaluated using the Area Under the Receiver Operating Characteristic (ROC) Curve (AUC), which provides a comprehensive measure of model accuracy across different classification thresholds. Furthermore, we ranked urinary microbiome according to Gini importance values.

### Association analysis for microbial taxa

Considering the power of GWAS tests, we refined our approach to focus on microbial taxa exhibiting occurrence rates exceeding 10% (present in at least 157 individuals) and an average relative abundance above 1×10^-4^. After filtering, the represented genera of these microbial taxa covered 99.73% of the whole community in the cohort. Subsequently, a total of 106 microbial genera and species were included for association analyses.

We examined the relationships between host genetics and the urine microbiome using two different statistical models: a linear model based on the relative abundance and a logistic model based on the presence/absence (P/A) of microbial features. For those taxa present in over 50% individuals, we used a linear model and transformed the relative abundance data using a natural logarithm. Subsequently, we calculated the residuals by employing the ’lm’ function in R: (log10(Microbe abundance) _ age + sex + BMI + sequencing read counts + top ten PCs). These residuals were then extracted using the residuals() function from the R stats package and applied in a univariate linear model for association analysis with genotypes. For the microbial taxa observed in more than 10% but less than 90% of individuals, we converted the data into presence/absence patterns to avoid issues with zero inflation. By dichotomizing these data, we could treat the bacterial abundance as a binary trait for logistic regression analysis. This analysis was adjusted for the same set of covariates as mentioned earlier. This dual approach allowed us to comprehensively assess the influence of host genetics on different aspects of the urine microbiome.

We searched the oral microbiome-related SNPs in the summary statistics data from this cohort as previously reported to examine their associations with host traits.

Additionally, we conducted GWAS association analyses similar to the steps described above for α-, β-diversity, and urinary clusters, however, no genome-wide significant signals were observed (**Supplementary Tables 13 & 14**). We also performed gene-environment interaction analyses on the top two significant loci, chr8:109442613 and chr2:191850055. These analyses, focusing on interactions with gender and serum testosterone levels, were executed using the --interaction parameter in PLINK.

### Two-sample MR analysis

To explore the causal relationships between the urinary microbiome and various diseases, we accessed summary statistics data of 42 diseases from Biobank Japan^40^. The two-sample bidirectional MR analysis was performed by applying the GSMR method^67^. Genetic variants with P < 1 × 10^−5^ and LD r^2^ <0.1 were also selected as instrumental variables for phenotypes in the Japan Biobank study.

### Data availability

The metagenomic sequencing data after removing host reads have been deposited to the CNGB Nucleotide Sequence Archive (CNSA: https://db.cngb.org/cnsa; accession number CNP0003862). The summary statistics that support the findings of this study including the associations between host genetics and urinary microbiome are available from https://db.cngb.org/search/project/CNP0005626/. According to the Human Genetic Resources Administration of China regulation and the institutional review board of BGI-Shenzhen related to protecting individual privacy, sequencing data are controlled-access and are available via application on request.

## Acknowledgments

We sincerely thank the support provided by China National GeneBank. We thank all the volunteers for their time and for self-collecting the oral samples using our kit. This work was supported by the National Natural Science Foundation of China (No. 32200548).

## Author contributions

X.L. and T.Z. conceived and organized this study. J.W. initiated the overall health project. X.X., X.J., L.X., Y.H., and Y.Z. contributed to organization of the cohort, the sample collection and questionnaire collection. H.L led the DNA extraction and sequencing. L.Z. and Z.Z. processed the metagenome data. X.L and X.T. processed the whole genome data. L.Z. and X.L performed the metagenome-genome-wide association analyses. L.Z., Z.Z., and X.L wrote the raw manuscript. All authors contributed to data and texts in this manuscript.

