## supplemental materials for "Unraveling the impact of host genetics and factors on the urinary microbiome in a young population"

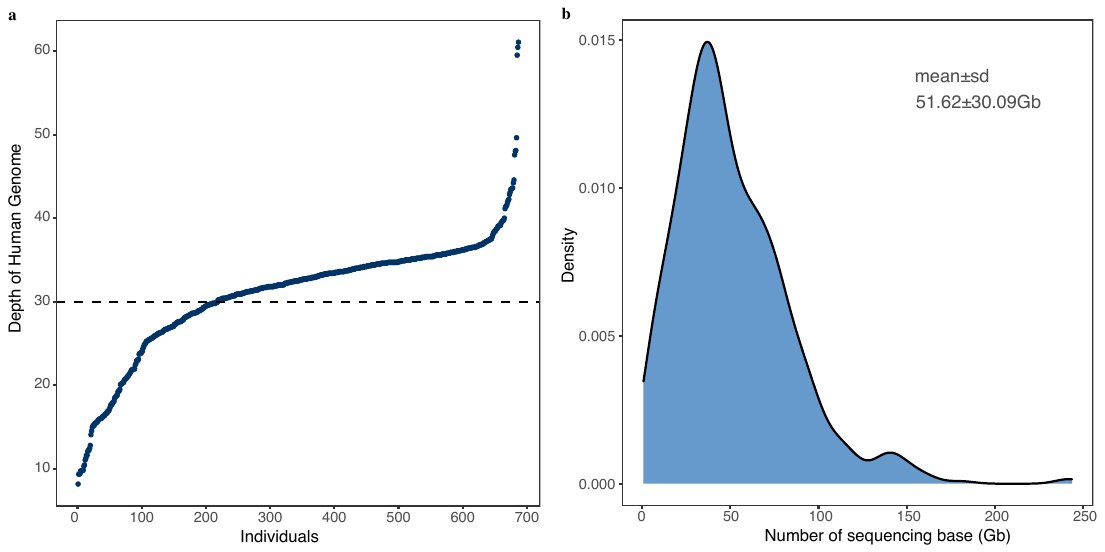

**Supplementary Fig. 1. Host genome and metagenome sequencing data production.**

(a) Depth distribution of 687 host genomes by integrating host whole genome sequencing data from blood sample and human-derived reads data extracted from urine sample. The mean depth is 30× (ranging from 8× to 61×). (b) Urine metagenome sequencing at an average of 51.62 ± 30.09 Gb after trimming low quality reads.

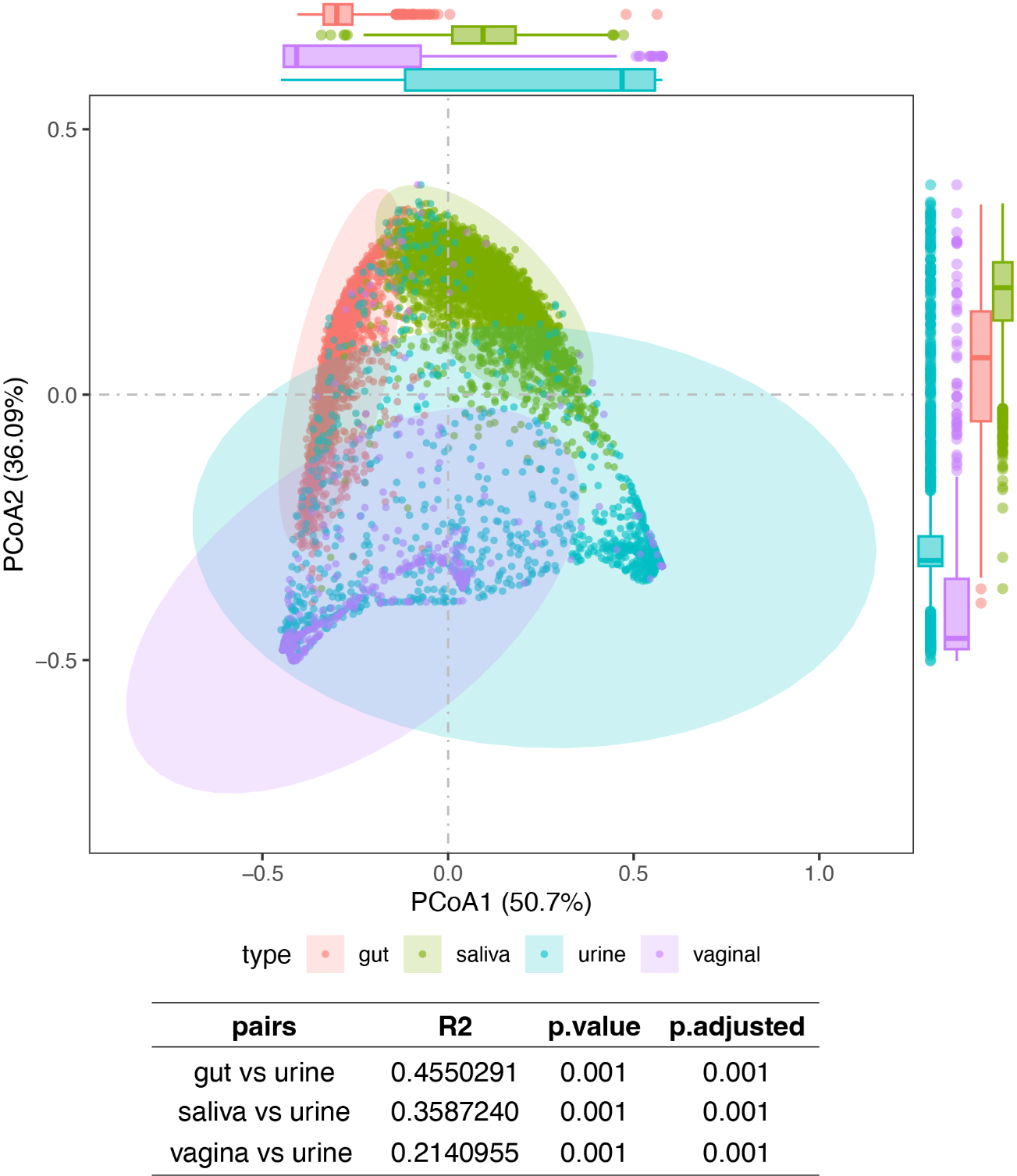

**Supplementary Fig. 2. Distribution of total samples’ PCOA at four sites: gut, salivary, urinary, and vaginal at phylum level.**

These included 3,222 saliva samples, 1,754 intestinal samples, 686 reproductive tract samples, and 1,579 urine samples. A PERMANOVA test was conducted to calculate the differences between each pairwise combination of these sample types.

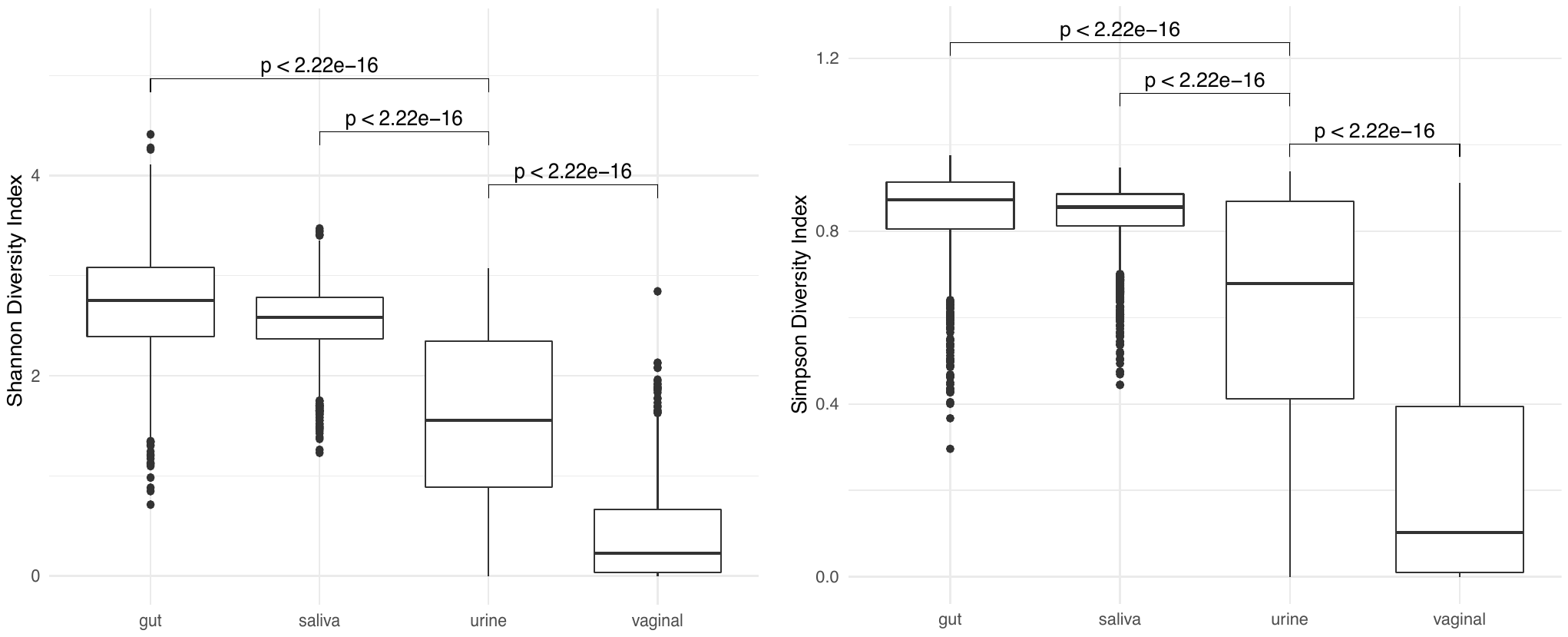

**Supplementary Fig. 3. Box plot of Shannon and Simpson Index distribution across four body sites: gut, saliva, vaginal, and urine.**

Shannon and Simpson indices were calculated based on species-level abundance data using the 'diversity' within the R package 'vegan'. The compared p-values of Wilcoxon test were also shown.

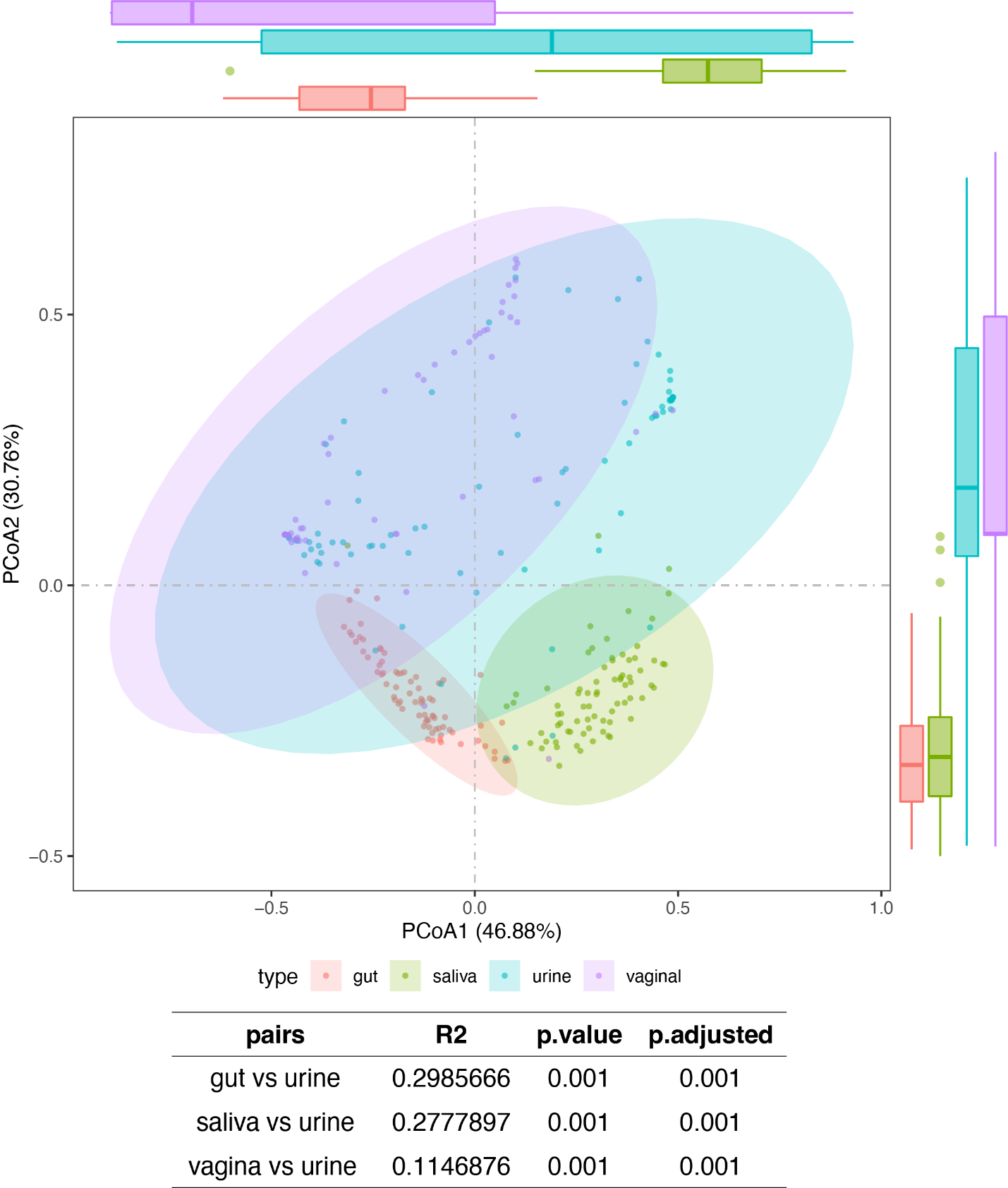

**Supplementary Fig. 4. Distribution of 75 collective samples’ PCOA at four sites: gut, salivary, urinary, and vaginal at phylum level.**

β-diversity was calculated based on species-level abundance data using the 'vegdist' functions within the R package 'vegan'. A PERMANOVA test was conducted to calculate the differences between each pairwise combination of these sample types.

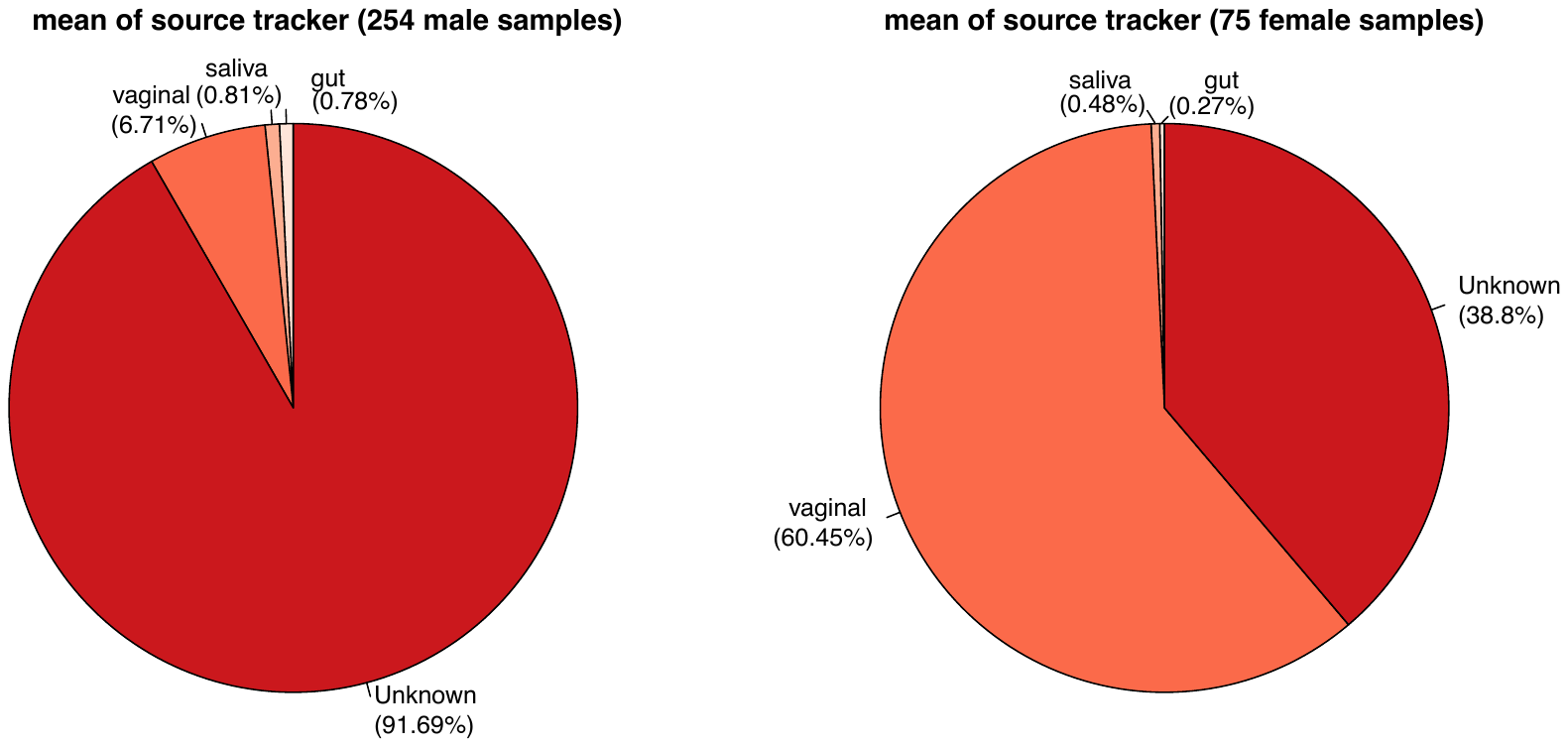

**Supplementary Fig. 5.** **Urinary microbiota source tracking in 75 samples** **with all four body sites.** Values represent the average percentage contribution of each source to the urinary microbiome.

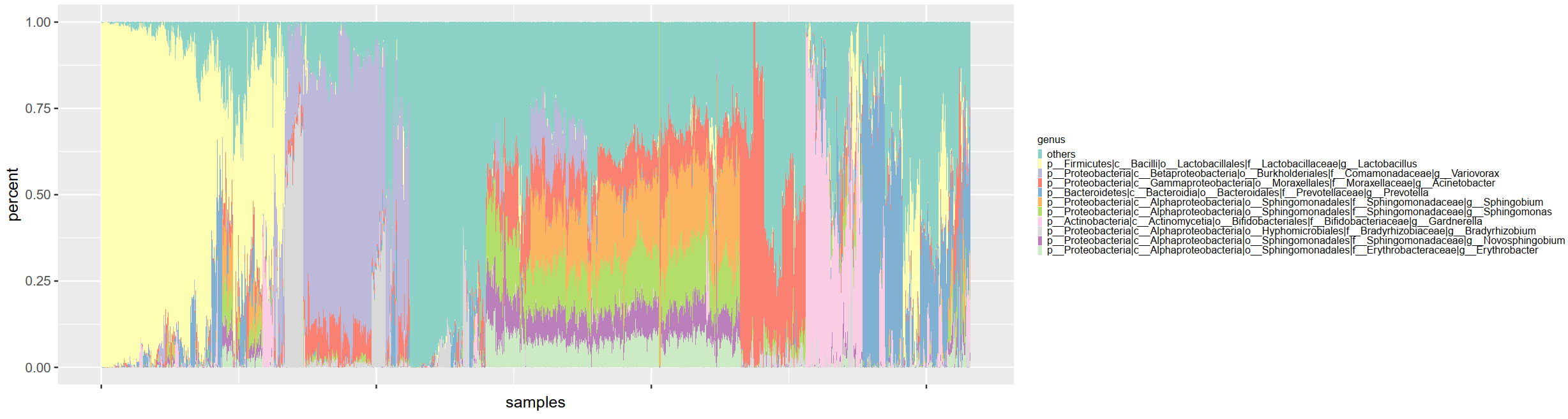

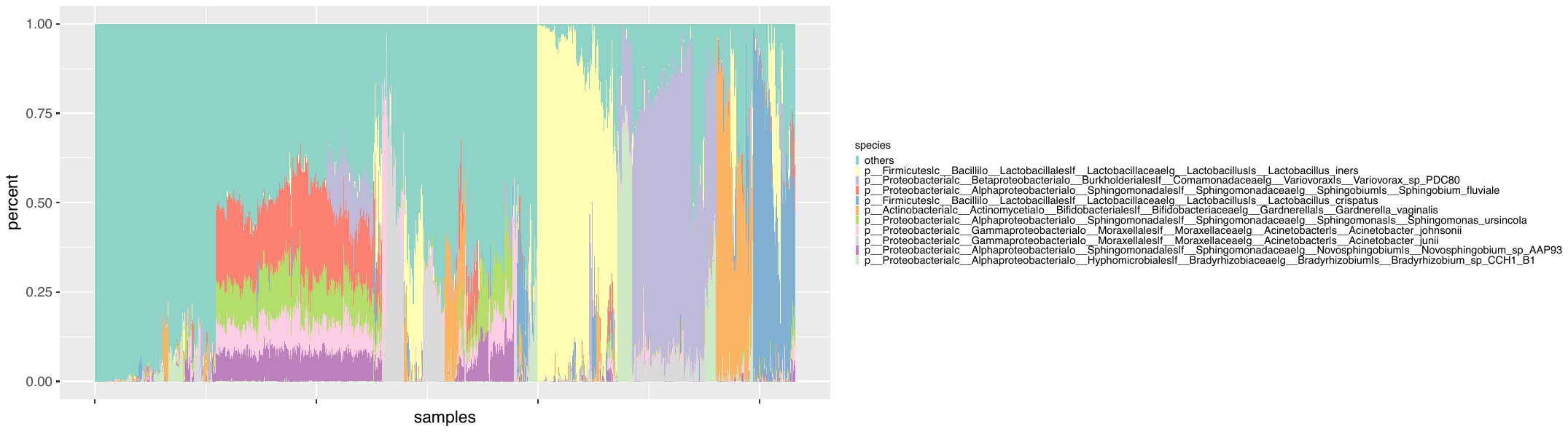

**Supplementary Fig. 6. Top 10 urinary bacteria with the highest relative abundance at both the genus and species levels in 1579 samples.**

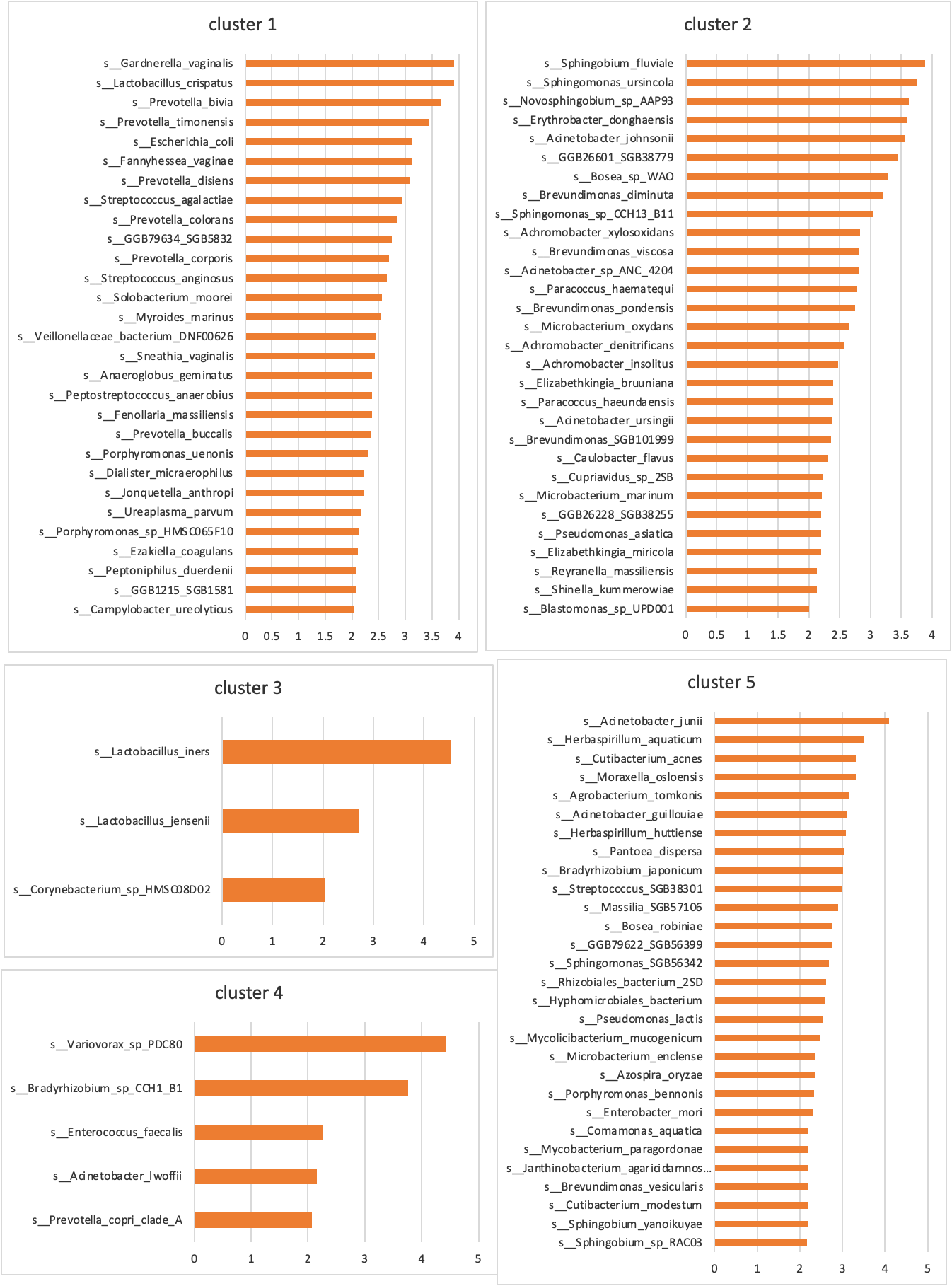

**Supplementary Fig. 7. The microbial species significantly enriched in five clusters respectively by ranking LDA scores from high to low.**

LDA scores were calculated by the Linear Discriminant Analysis Effect Size method.

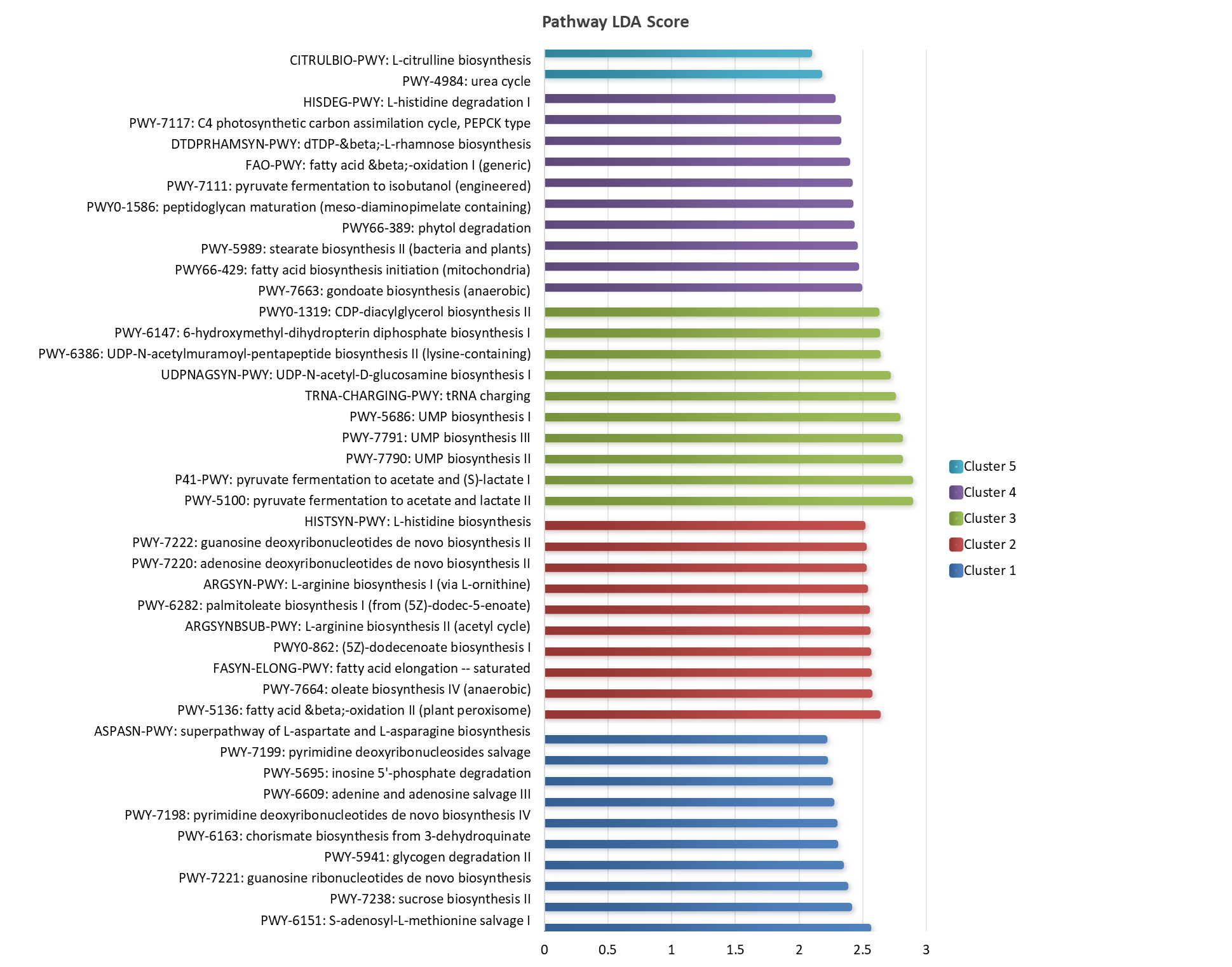

**Supplementary Fig. 8. The top 10 microbial pathways significantly enriched in five clusters respectively by ranking LDA scores from high to low.**

LDA scores were calculated by the Linear Discriminant Analysis Effect Size method.

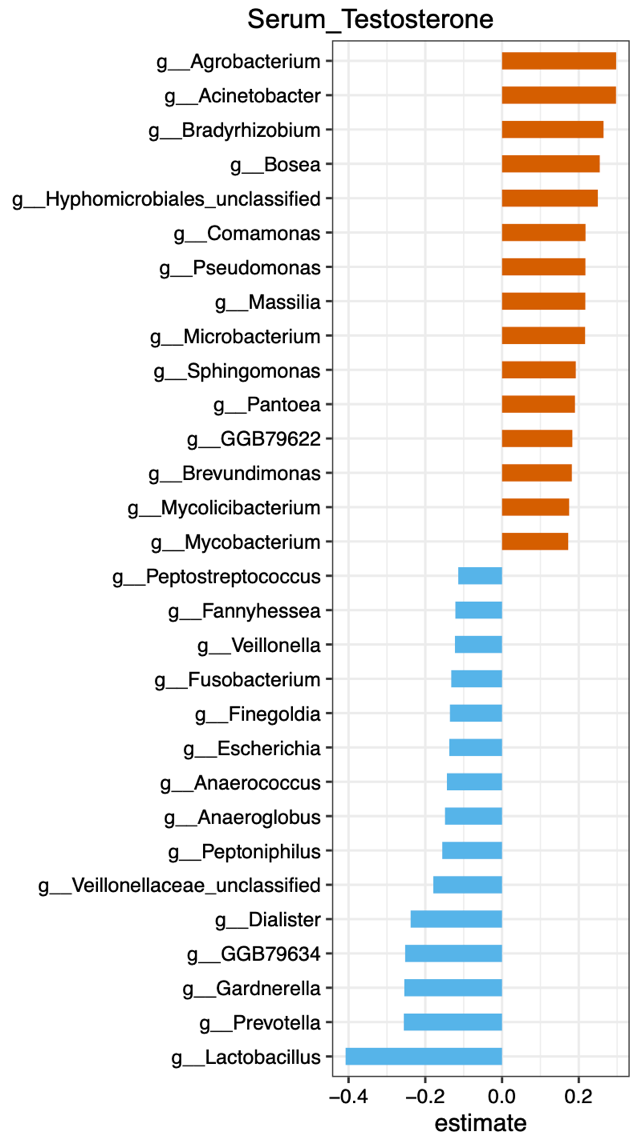

**Supplementary Fig. 9. The top 15 positive and top 15 negative correlates of serum testosterone in explaining urinary microbiota.**

We employed Bray-Curtis distance-based redundancy analysis (dbRDA) and determined the variance explained by these factors using the 'capscale' function in the vegan package.

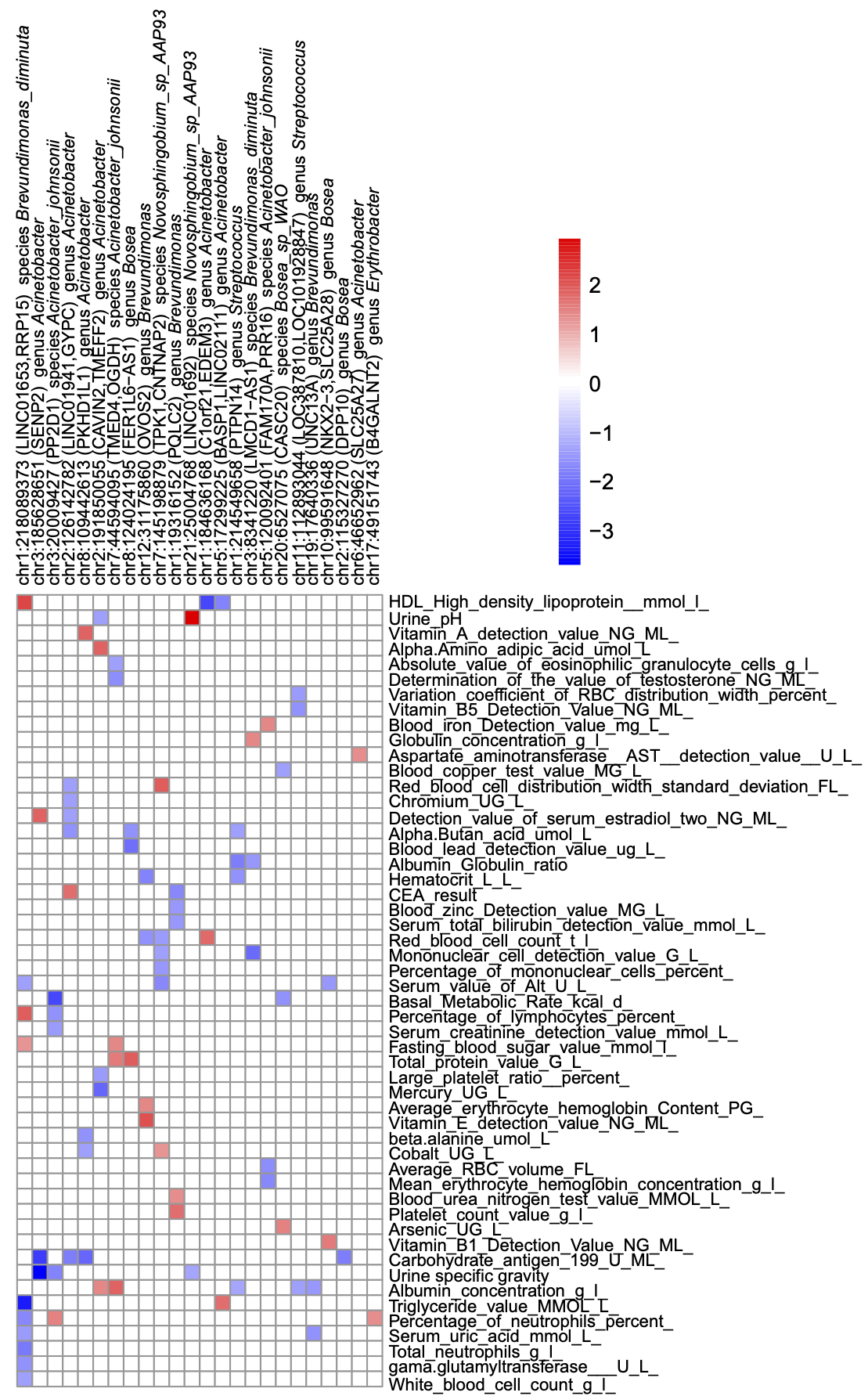

**Supplementary Fig. 10. The correlation between top loci and our 2018 metabolic database.** The values in the heatmap are derived from the 'p_adjusted' column in Supplementary table 10.

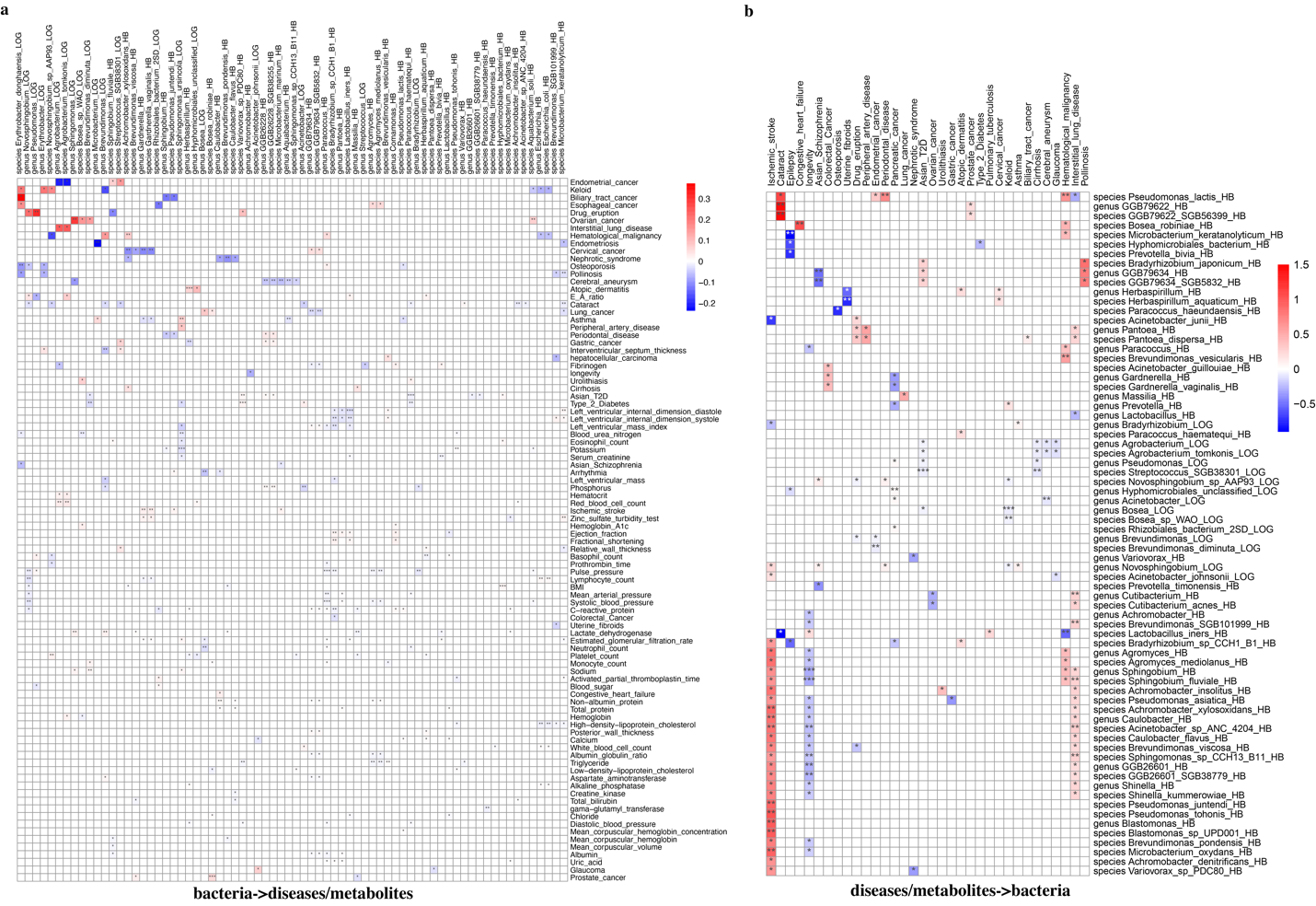

**Supplementary Fig. 11. Two-sample MR identifying significant causal relationships urinary bacterium and disease from BBJ**.

The numerical values in the heatmap are sourced from the 'bxy' column in Supplementary Table 11. The notation of *** indicates p < 0.001, ** indicates p < 0.01, and * indicates p < 0.05. a) Exposure: bacteria, Outcome: diseases/metabolites; b) Exposure: diseases/metabolites, Outcome: bacteria.

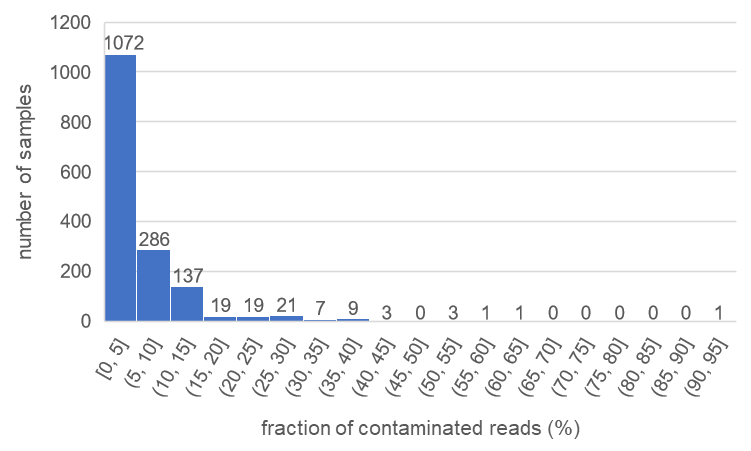

**Supplementary Fig. 12. Distribution of urinary microbiota contamination rates in 1579 samples**

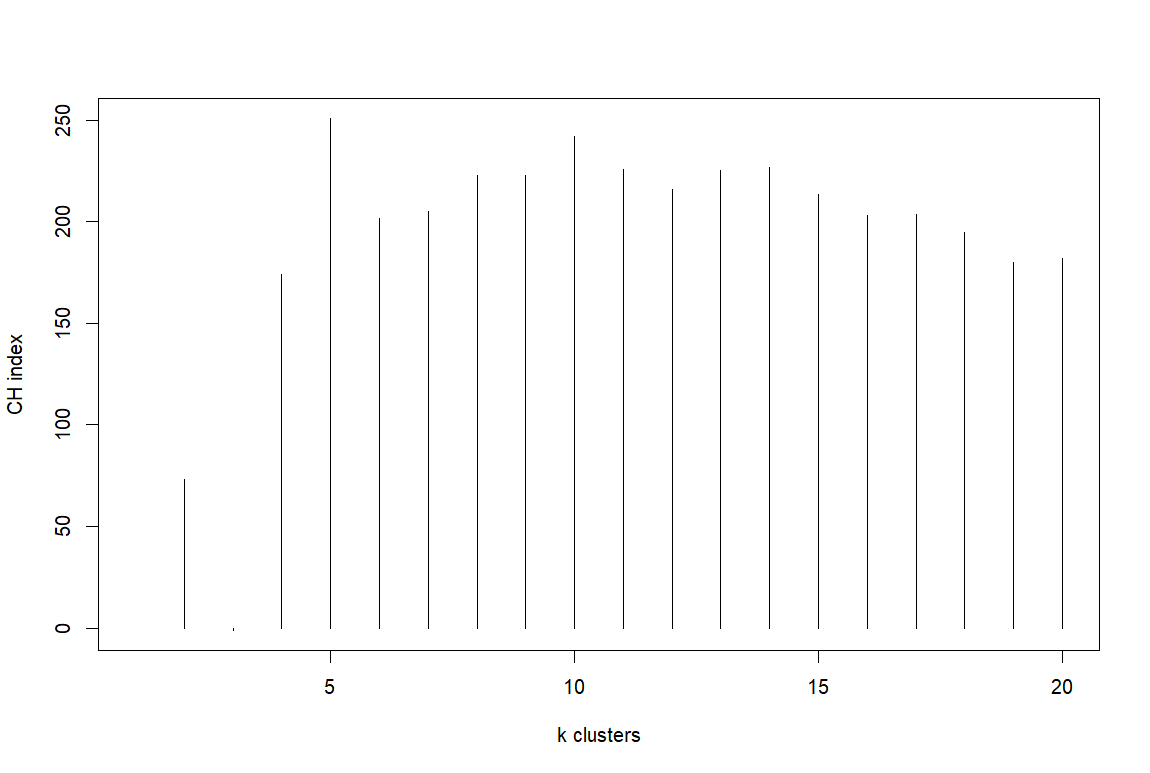

**Supplementary Fig.13. Calinski-Harabasz (CH) Index to determine the optimal number k of urinary clusters.**

Using Partitioning Around Medoids (PAM) clustering algorithm to calculate CH values under different numbers of clusters.
